# Evaluating cacao pod resistance to black pod disease caused by three *Phytophthora* species

**DOI:** 10.64898/2026.09.17.752234

**Authors:** Mariana Herrera-Corzo, Ina Schlathoelter, Adriana Arciniegas-Leal, Bryan A. Bailey, Alina S. Puig, Yeirme Yaneth Jaimes Suárez, Shamala Sundram, Thi Thu Nga Nguyen, Dahyana Santos Britto, Dayana C. Rodezno, Derek R. Drost, Jean-Phillipe Marelli, Jeremy T. Brawner, Erica M. Goss

## Abstract

Black pod disease is a major threat to global cacao (*Theobroma cacao* L.) production. It is caused by multiple *Phytophthora* species with geographic ranges from regionally restricted to pantropical, and developing resistant cultivars requires screening against the diversity of these pathogens. This study evaluated the response of 25 cacao clones to 13 isolates of *P. palmivora* (eight), *P. megakarya* (two), and *P. theobromicola* (three) by nonwounding inoculation of detached pods. Lesion size and lesion expansion status were evaluated 4 days after inoculation. The linear mixed model included 178 combinations among 17 clones and 12 isolates, each represented by at least two biological replicate pods. Lesion size differed among clones (F_(16, 449.2)_ = 25.16) and among isolates (F_(11, 416.0)_ = 32.90), and the clone by isolate interaction was significant (F_(150, 465.4)_ = 1.72; all *P* < 0.001). PENTAGONA-17 had the smallest estimated marginal lesion size (9.3 mm) and fitted values of 3.8 to 16.9 mm across 11 isolates representing all three species. ARF-31, ICS-47, POUND-7, and PMCT-73 had the next four smallest estimates, whereas NA-246 had the largest (36.7 mm). Isolate estimates varied more than fourfold, from 9.6 to 42.0 mm. Despite a marginal estimate of 12.9 mm, POUND-7 had a fitted estimate of 49.2 mm with *P. megakarya* GH34. The probability of lesion expansion also differed among clones and isolates (both *P* < 0.001). Together, these findings identify candidates for further resistance evaluation and show that multi-isolate screening provides a stronger basis for selecting materials for black pod resistance breeding.

## Introduction

Cacao (*Theobroma cacao* L.) supports a global chocolate industry valued at approximately US $140 billion and provides livelihoods for an estimated 5 to 6 million predominantly smallholder farmers worldwide (Boysen et al. 2023). Black pod disease, caused by multiple *Phytophthora* species, is one of the most important constraints to cacao production worldwide, with annual losses estimated at approximately 0.87 million metric tons of dried cacao beans, representing about US$2 billion (Marelli et al. 2019; Morales-Cruz et al. 2020). *P. palmivora* is globally distributed and can infect multiple cacao tissues, causing annual yield losses estimated around 20-30%, while *P. megakarya* is restricted to West and Central Africa. It can cause crop losses of up to 90% in the absence of effective management (Ali et al. 2016; Guest 2007; Morales-Cruz et al. 2020). More recently, *P. theobromicola* was described as a causal agent of black pod disease in Brazil (Decloquement et al. 2021). The variation in geographic distribution and disease severity among *Phytophthora* species underscores the complexity of black pod disease and the ongoing challenges these pathogens present to cacao production.

Breeding for resistance is the most sustainable long-term strategy for managing black pod disease, and several sources of resistance or reduced susceptibility to *Phytophthora* spp. have been identified: SCA-6, is widely used as a source of reduced susceptibility to *P. palmivora* and *P. megakarya* (Nyassé et al. 2007); POUND-7 demonstrates resistance to *P. palmivora* but is susceptible to frosty pod rot caused by *Moniliophthora roreri* (Baruah et al. 2024; Brown et al. 2007; Gutiérrez et al. 2021), whereas UF-273 (Type I) is resistant to frosty pod rot but susceptibility to black pod disease (Brown et al. 2007; Gutiérrez et al. 2021). Complete resistance to *P. megakarya* has not yet been found, although screening studies have revealed significant variations in resistance among cacao genotypes. (Nyassé et al. 2007; Ofori et al. 2023). Recent screening against a diverse collection of *P. megakarya* isolates has shown a similar pattern (I. Schlathoelter et al. unpublished data). Upper Amazon clones such as SCA-6, POUND 7, PA 150, and T85/799 show comparatively low susceptibility and are valuable parents for breeding (Nyassé et al. 2007). While wild cacao populations exhibit substantial genetic diversity, cacao breeding has historically relied on a limited subset of that diversity, potentially restricting the range of resistance alleles available for improvement (Motamayor et al. 2008; Zhang and Motilal 2016). Cacao breeding is also constrained by the species’ highly outcrossing nature and long generation time. Modern cacao breeding programs have existed for less than a century and have advanced only a few generations, with breeding historically relying heavily on clonal selection and a limited number of crossing cycles (DuVal et al. 2017; Rodriguez-Medina et al. 2019; Zhang and Motilal 2016). Broadening the genetic base and developing new varieties with enhanced, durable resistance are priorities for cacao breeding (Nyassé et al. 2007; Zhang and Motilal 2016).

Evaluating cacao responses to individual *Phytophthora* species provides useful breeding information; identifying germplasm with resistance across species is essential. Nyassé et al. (2007) reported relatively consistent resistance rankings across West African field trials where different *Phytophthora* species predominated. Similarly, controlled evaluations of cacao progenies against *P. palmivora* and *P. megakarya* showed consistent relative resistance rankings across the two species (Nyadanu et al. 2012b). However, consistent resistance across species is not always observed; among 262 F1 progeny from a TSH x CCN 51 cross evaluated against *P. palmivora, P. capsici,* and *P. citrophthora*, only ten were resistant to all three species (Barreto et al. 2015). Variation within species adds another level of complexity. For example, *P. palmivora* isolates from different cacao-growing regions in Colombia differed significantly in aggressiveness on cacao pods (Rodríguez Polanco et al. 2022). Differences in aggressiveness have also been reported among *P. theobromicola* isolates on cacao pods (Decloquement et al. 2021). Together, these findings highlight the importance of evaluating cacao resistance across multiple *Phytophthora* species and isolates. The variation in resistance across *Phytophthora* species and isolates reflects a genetically complex host response. Cacao resistance is quantitatively inherited and controlled by multiple loci distributed across the genome (Mucherino Muñoz et al. 2021). Some loci have been associated with resistance to more than one *Phytophthora* species, whereas others are species-specific, and the degree of overlap differs among studies (Barreto et al. 2018; Risterucci et al. 2003). At the molecular level, resistant and susceptible cacao genotypes also differ in the expression and polymorphism of defense-related genes (Pokou et al. 2019).

Beyond host variation, genetic differences are also evident among *Phytophthora* species and populations. Comparative genomic analyses of *P. palmivora* and *P. megakarya* revealed differences in genome architecture and expansions of gene families associated with infection (Morales-Cruz et al. 2020). Intraspecific genomic variation is also evident in *P. theobromicola*, including structural variation and variable effector repertoires among isolates (García et al. 2026). More recently, high genotypic diversity was reported among *P. palmivora* isolates of cacao in Indonesia (Brugman et al. 2022).

Given this variation in both cacao and its *Phytophthora* pathogens, phenotypic assays remain crucial for evaluating resistance, although the response measured can depend on the plant tissue and the experimental design. Leaf-based assays are practical for early screening before pod production, but results obtained from leaves do not always correspond closely with pod or field responses, and can be influenced by leaf developmental stage and experimental conditions (Iwaro et al. 1997a; Nyassé et al. 1995; Tahi et al. 2006). Because black pod disease refers to the pod-rot phase that directly reduces harvestable yield, pod inoculation allows evaluation of disease response in the organ most relevant to the disease outcome. Detached pod assays also allow inoculum concentration, incubation conditions, and assessment timing to be standardized, improving the repeatability of the resistance evaluations and allowing cacao genotypes to be compared under a common experimental framework (Iwaro et al. 2005; Nyadanu et al. 2012b). These assays complement rather than replace field evaluations; detached pod responses have shown significant associations with field disease incidence, whereas field responses are also influenced by environmental variation (Iwaro et al. 2005; Ofori et al. 2023).

Studies of cacao response to black pod disease have evaluated different combinations of *Phytophthora* species and isolates, but the three species targeted here have not been examined together in an evaluation using multiple isolates and pod inoculation.

Nyadanu et al. (2012) assessed cacao responses to *P. palmivora* and *P. megakarya* using leaf disc, detached pod, and field evaluations, whereas Barreto et al. (2015) evaluated *P. palmivora*, *P. capsici*, and *P. citrophthora* using one isolate of each species in a leaf disc assay. *P. theobromicola* has been characterized in pathogenicity tests on pods in four cacao clones (Decloquement et al. 2021), but it has not been included in the broader multi-species resistance evaluations described above. Dadzie et al. (2026) recently evaluated 28 cacao genotypes using detached pod inoculation with a single *P.* palmivora isolate.

In this study, we evaluated cacao response to black pod disease using a standardized detached pod inoculation protocol. The screening included 25 cacao clones and 13 isolates representing *P. palmivora* (eight isolates), *P. megakarya* (two isolates), and *P. theobromicola* (three isolates). Isolates were selected to include different geographic origins and within-species variation available in our *Phytophthora* collection, while considering the known distribution of each species. Our objectives were to i) identify cacao genotypes showing relatively low lesion development across the *Phytophthora* species, and isolates tested, ii) compare aggressiveness among isolates evaluated within each species, and iii) assess whether cacao responses to individual isolates differed among genotypes by testing the genotype by isolate interaction in lesion development. This comparative evaluation of cacao response to black pod disease across species and isolates provides phenotypic information relevant to resistance breeding.

## Materials and Methods

### Cacao clone and pathogen isolate selection

Cacao pods were obtained from the International Cacao Collection at Tropical Agricultural Research and Higher Education Center (CATIE *Centro Agronómico Tropical de Investigación y Enseñanza*) in Turrialba, Costa Rica, located at 604 meters above sea level, with an average temperature of 22.5°C, and annual precipitation of 2645 mm (Mata-Quirós et al. 2017). A total of 25 cacao clones were selected to span a range of previously characterized responses to *P. palmivora* and *P. megakarya.* Selection was informed by unpublished CATIE screening data for pod responses to *P. palmivora* isolate GH49, an isolate previously characterized by Ali et al. (2016) and by controlled screening of the CATIE germplasm collection with *P. megakarya* isolates (I. Schlathoelter et al. unpublished data) (Table 1). Healthy, uniform cacao pods obtained by hand pollination were harvested at approximately four months of age and shipped to the Plant Pathology Department at the University of Florida (Gainesville, Florida, USA) under USDA APHIS PPQ permits for cacao pod importation 556-20-330-01055 and 556-23-242-14283.

**Table 1.** Characteristics and observed screening responses of the 25 cacao clones evaluated in this study.

| Cacao clone | Origin<sup>a</sup> | Principal<br>genetic group<sup>b</sup> | Documented<br>pedigree<sup>c</sup> | Isolates<br>evaluated<br>(n) | Qualifying<br>combinations<br>(n)<sup>d</sup> | Replicate<br>pods per<br>isolate<br>(min–max) | Observed<br>lesion size,<br>mean (range),<br>mm<sup>e</sup> |
| --- | --- | --- | --- | --- | --- | --- | --- |
| **PENTAGONA-17** | Nicaragua | Not represented | — | 11 | 11 | 2–5 | 10.3 (0–38) |
| **PNG-197** | Papua New<br>Guinea | Amelonado-Hybrid | — | 3 | 0<sup>d</sup> | 1–1 | 10.7 (6–14) |
| **POUND-19-A** | Peru | Nanay-Hybrid | — | 7 | 4<sup>d</sup> | 1–2 | 11.8 (0–55) |
| **ARF-31** | Costa Rica | Contamana-Hybrid | SCA-6 × CC-42 | 10 | 7 | 1–4 | 14.0 (0–50) |
| **PMCT-73** | Costa Rica | Contamana-Hybrid | POUND-7 × UF-667 | 11 | 11 | 3–5 | 16.5 (0–74) |
| **POUND-7** | Peru | Nanay | — | 13 | 12 | 1–7 | 16.7 (0–64) |
| **HY-2714182** | Trinidad and<br>Tobago | Iquitos-Hybrid | IMC-67 × PA-218 | 4 | 2<sup>d</sup> | 1–2 | 17.2 (0–56) |
| **P-19** | Peru | Nanay-Hybrid | — | 12 | 12 | 2–8 | 17.4 (0–57) |
| **ICS-47** | Trinidad | Amelonado-Hybrid | Criollo hybrid | 12 | 10 | 1–4 | 18.0 (0–58) |
| **PMCT-37** | Costa Rica | Amelonado-Hybrid | UF-676 ×<br>POUND-7 | 12 | 11 | 1–4 | 18.8 (0–60) |
| **COCA-3370-5** | Ecuador | Iquitos-Hybrid | — | 9 | 2<sup>d</sup> | 1–2 | 19.2 (4–48) |
| **ARF-10** | Belize | Maranon-Hybrid | — | 4 | 0<sup>d</sup> | 1–1 | 23.1 (0–49) |
| **CCN-51-T1** | Ecuador | Amelonado-Hybrid | (IMC-67 × ICS-95) × Oriente 1 | 12 | 12 | 2–5 | 23.7 (0–68) |
| **EEG-65** | Brazil | Amelonado | — | 12 | 10 | 1–3 | 26.7 (2–70) |
| **CRIOLLO-34** | Nicaragua | Amelonado-Hybrid | — | 12 | 11 | 1–6 | 26.7 (0–64) |
| **LCTEEN-37-F** | Ecuador | Curaray-Hybrid | — | 8 | 0<sup>d</sup> | 1–1 | 27.5 (5–52) |
| **SCA-6** | Peru | Not represented | — | 9 | 4<sup>d</sup> | 1–2 | 28.2 (0–52) |
| **CATIE-1000** | Costa Rica | Amelonado-Hybrid | POUND-12 ×<br>CATONGO | 12 | 12 | 2–5 | 28.6 (0–60) |
| **UF-273-T1** | Costa Rica | Nacional-Hybrid | — | 13 | 11 | 1–4 | 29.7 (0–66) |
| **PA-150** | Peru | Iquitos-Hybrid | — | 11 | 11 | 2–4 | 31.6 (0–62) |
| **IMC-60** | Peru | Iquitos | — | 6 | 6 | 2–2 | 33.3 (0–61) |
| **PA-4** | Peru | Maranon | — | 10 | 8 | 1–4 | 33.5 (0–63) |
| **NA-246** | Peru | Maranon | — | 12 | 12 | 2–6 | 39.2 (0–70) |
| **CCN-10** | Ecuador | Amelonado-Hybrid | — | 13 | 12 | 1–6 | 39.7 (0–82) |
| **PMCT-31** | Costa Rica | Amelonado-Hybrid | Open-pollinated<br>selection | 3 | 3<sup>d</sup> | 3–4 | 48.5 (0–58) |

Clones were genotyped for a companion study with a 10,000-position single nucleotide polymorphism panel, and genetic group membership was inferred with STRUCTURE (Pritchard et al. 2000) against the reference groups of Motamayor et al. (2008) and Zhang et al. (2012) (I. Schlathoelter, unpublished data; Table 1). Documented pedigree and provenance were compiled from CATIE germplasm records, and parentage was reported only when explicitly documented (Table 1).

Thirteen *Phytophthora* isolates representing three species were selected for evaluation: eight *P. palmivora*, two *P. megakarya*, and three *P. theobromicola*. The isolate panel included multiple geographic origins available in our collection. Species identity, geographic origin, and collection year, when available, are provided in Table 2. All isolates were handled following the standard operating procedures associated with corresponding USDA-APHIS-PPQ permits, P526-21-01779 and 526-24-197-95249.

**Table 2.** *Phytophthora* isolates evaluated in this study.

| Isolate | Species | Geographic<br>origin | Host<sup>a</sup> | Collection<br>year | Clones<br>evaluated<br>(n) | Qualifying<br>combinations<br>(n)<sup>c</sup> | Replicate<br>pods per<br>clone (min–<br>max) | Observed<br>lesion size,<br>mean (range),<br>mm<sup>b</sup> |
| --- | --- | --- | --- | --- | --- | --- | --- | --- |
| CATP5 | *P. palmivora* | Costa Rica | Theobroma<br>cacao | 2022 | 20 | 16 | 1–6 | 24.9 (0–66) |
| GH49<sup>d</sup> | *P. palmivora* | Ghana | Theobroma<br>cacao | 2013 | 25 | 21 | 1–7 | 14.4 (0–51) |
| KORO | *P. palmivora* | Papua New<br>Guinea | Theobroma<br>cacao | — | 19 | 16 | 1–6 | 19.3 (0–52) |
| MCCS-MB-01919 | *P. palmivora* | Brazil | Theobroma<br>cacao | — | 17 | 14 | 1–6 | 27.5 (0–62) |
| MCCS-MB-01994 | *P. palmivora* | Brazil | Theobroma<br>cacao | — | 19 | 16 | 1–6 | 26.3 (0–68) |
| NPP3 | *P. palmivora* | Colombia | Theobroma<br>cacao | — | 20 | 17 | 1–6 | 34.7 (0–66) |
| PP11 | *P. palmivora* | Malaysia | Theobroma<br>cacao | 2014 | 17 | 15 | 1–6 | 26.2 (0–56) |
| Phy-VL | *P. palmivora* | Vietnam | Theobroma<br>cacao | — | 19 | 15 | 1–6 | 12.8 (0–63) |
| GH34<sup>d</sup> | *P. megakarya* | Ghana | Theobroma<br>cacao | 2013 | 24 | 21 | 1–6 | 43.9 (0–82) |
| ZTH0466 | *P. megakarya* | Cameroon | Theobroma<br>cacao | — | 19 | 17 | 1–6 | 21.2 (0–56) |
| MCCS-MB-01934 | *P. theobromicola* | Brazil | Theobroma<br>cacao | 2017 | 14 | 9 | 1–4 | 31.1 (0–72) |
| MCCS-MB-01952 | *P. theobromicola* | Brazil | Theobroma<br>cacao | 2017 | 8 | 1<sup>c</sup> | 1–2 | 33.4 (5–70) |
| MCCS-MB-01960 | *P. theobromicola* | Brazil | Theobroma<br>cacao | 2017 | 20 | 16 | 1–8 | 20.3 (0–64) |

by isolate combinations that qualified for the inferential analyses; a combination qualified when it had at least 12 site records after imputation, equivalent to at least two replicate pods. The value marked <sup>c</sup> is below five; this isolate was excluded from the inferential analyses (retained isolates had at least nine qualifying combinations). <sup>d</sup> GH49 and GH34 are the isolates reported as Gh-ER1349 (*P. palmivora*) and Gh-ER1334 (*P. megakarya*) by Ali et al. (2016) and sequenced as Pp2 and Pm1 by Morales-Cruz et al. (2020). n = number.

### Isolate maintenance and Inoculum preparation

Isolates were maintained on 10% clarified tomato medium (CTM; clarified tomato juice with 1g calcium carbonate per100 mL), solidified with 1.5% agar, amended with Delvocid (50% Pimaricin) 10 mg/L, Ampicillin 250 mg/L, and Rifamycin-SV 10 mg/L (Ferguson and Jeffers 1999). To induce sporangia formation, two mycelial plugs (5 mm diameter) from the actively growing margins of 7-day-old colonies were transferred to fresh 10% CTM without antibiotics and incubated at 25°C in the darkness for five days, followed by exposure to fluorescent light ( 24 watts, 5000 lumens, cool white) for an additional five days at 25°C (Puig et al. 2021).

Zoospores were released by flooding each plate with 15 mL of chilled (4 °C) sterile distilled water. For *P. palmivora* and *P. megakarya*, plates were kept at 4°C for 20 minutes, then transferred to room temperature for 30 minutes (Ali et al. 2016). For *P. theobromicola*, plates were held at 4°C for 45 minutes, then incubated at 28°C for 30 minutes (Schmidt et al. 2023). A 2 mL aliquot of the suspension was transferred to a glass test tube using a wide-mouth pipette tip and vortexed for 2 minutes to encyst the zoospores before counting with a Neubauer hemocytometer. The concentration measured in the encysted aliquot was then used to dilute the remaining motile suspension with sterile distilled water to 50,000 zoospores/mL for inoculation. Each suspension was used within 1 to 2 hours, and its concentration was reconfirmed by hemocytometer at the end of each inoculation session.

### Detached pod zoospore patch assay

Pod response to each isolate was evaluated using the Detached Pod-Zoospores Patch (DP-ZP) assay described by Gitto et al.(2026), adapted from Phillips-Mora and Galindo (1989). Approximately 10 (±1) days elapsed between pod harvest and inoculation.

Before inoculation, pods were inspected, rinsed with reverse-osmosis water, and labeled with clone and isolate information; pods that arrived in poor condition or showed deterioration were excluded. Six inoculation sites were distributed evenly around each pod, in the furrows between pod ridges, and at least 25 mm from the stem end, the pod tip, and adjacent inoculation sites. The inoculation was performed while maintaining pod surface integrity (non-wounding). A 1 cm^2^ tissue patch was placed at each site, and 30 μL of zoospore suspension at 50,000 zoospore/mL was pipetted onto each patch, delivering 1,500 zoospores per inoculation site. No mock-inoculated control was included; responses were compared among clones and isolates rather than against an uninoculated baseline.

Inoculated pods were placed in sealed boxes lined with dampened paper towels to maintain relative humidity near 100% and held in two growth chambers (Percival Scientific, model I36NL) at 25°C in darkness. At 4 days after inoculation, each pod was inspected to determine whether the inoculation sites could be evaluated reliably. Sites were recorded as missing when lesion boundaries could not be distinguished because lesions from adjacent inoculation sites had coalesced. Pods were excluded and flagged for replacement when extensive background lesions, scarring, discoloration, or other tissue deterioration masked or overlapped the inoculation sites, preventing evaluation. For evaluable sites, reaction phenotype was first recorded qualitatively as 0, no visible response; 1, loc localized darkening or necrosis confined to the inoculation area without lesion expansion; 2, localized necrosis followed by an expanding lesion extending beyond the inoculation area; or 3, a typical expanding lesion. These categories describe the observed phenotype and do not imply a specific defense mechanism. Lesion development was then quantified at 4 days after inoculation by measuring the maximum vertical and horizontal dimensions of the visible lesion to the nearest millimeter and averaging the two measurements to obtain lesion size. Sites with no visible lesion were recorded as 0 mm. When localized necrotic flecking was accompanied by lesion expansion, the dimensions of the expanding lesion were measured. Each pod was photographed before inoculation and again after evaluation at 4 days after inoculation for visual documentation.

### Experimental design and data processing

The experiment was designed to evaluate the pod responses of 25 cacao clones to 13 *Phytophthora* isolates, corresponding to 325 possible clone by isolate combinations. Each combination targeted three independent biological replicate pods, with six inoculation sites per pod, for a total of 18 site observations per combination. The six sites were treated as subsamples nested within their respective pod. To accommodate differences in flowering time and pod availability among clones, CATIE supplied 737 pods in ten shipments, each representing a separate bioassay. The reference isolates GH34 (*P. megakarya*) and GH49 (*P. palmivora*) were included in nine of the ten bioassays, providing repeated reference isolates across shipments. Variation in pod availability among cacao clones and exclusion of material unsuitable for evaluation resulted in an incomplete, unbalanced design with unequal replication among clone by isolate combinations. Observations from all 25 clones and 13 isolates were retained for descriptive summaries. Inferential analyses used the subset meeting the inclusion criteria described below. The coverage before filtering , and the retained combinations are shown in Supplementary Table S1.

We performed data processing in two stages. First, pods with fewer than three observed lesion size values were excluded. For retained pods, missing vertical and horizontal measurements were imputed separately using the corresponding mean for that dimension within the pod, and lesion size was recalculated. Overall, 265 of 4,416 site records (6.0%) required imputation. Second, clone by isolate combinations with at least 12 site records after imputation were retained. In the dataset evaluated at this stage, this criterion was equivalent to retaining combinations represented by at least two biological replicate pods. Clone and isolate levels represented by fewer than five qualifying combinations were subsequently excluded from inferential analyses. After applying these criteria, one isolate, MCCS-MB-01952 (*P. theobromicola*), and eight clones PNG-197, PMCT-31, ARF-10, HY-2714182, LCTEEN-37-F, SCA-6, COCA-3370-5, and POUND-19-A, were excluded from the inferential dataset. Their observations remained in the descriptive dataset summarized in Supplementary Table S1. The final lesion size dataset contained 3,918 site observations nested within 654 biological replicate pods across ten bioassays. It represented 17 clones and 12 isolates, with observations for 178 of the 204 possible combinations among these retained clones and isolates (87.3%; Supplementary Table S1, bold cells).

### Statistical Analysis

All statistical analyses were conducted using R v4.4.1 (R Core Team 2024). Linear and generalized linear mixed models were fitted using lme4 v1.1-37 (Bates et al., 2015).

Fixed effect F tests with Satterthwaite degrees of freedom and likelihood ratio tests of random effects used lmerTest v3.1-3 (Kuznetsova et al. 2017), whereas Type II Wald χ² tests for the binary model used car v3.1-3 (Fox and Weisberg 2019). Estimated marginal means and pairwise comparisons were obtained using emmeans v2.0.0 (Lenth and Piaskowski 2025), with multcomp v1.4-29 (Hothorn et al. 2008) and multcompView v0.1-10 (Graves et al. 2024) used for compact letter displays. Data manipulation, factor handling, descriptive statistics, and table formatting used dplyr v1.1.4 (Wickham et al. 2023), forcats v1.0.1 (Wickham 2025), psych v2.5.6 (Revelle 2025), and pander v0.6.6 (Daróczi and Tsegelskyi 2025), respectively. Figures were produced using ggplot2 v4.0.0 (Wickham 2016), ggbeeswarm v0.7.2 (Clarke et al. 2023), ggpubr v0.6.2 (Kassambara 2025) and patchwork v1.3.2 (Pedersen 2025), with EnvStats v3.1.0 (Millard 2013) used for sample size annotations. Analysis scripts and the underlying data are available at https://github.com/maherreracorzo.

Lesion size was the primary continuous response and was analyzed using a linear mixed model with cacao clone, *Phytophthora* isolate, and their interaction as fixed effects, representing comparisons among the selected experimental materials. Bioassay and individual pod were included as random intercepts. Each pod was uniquely identified by its clone, pod number, bioassay, and isolate. Inoculation sites were treated as subsamples nested within each pod. Lesion size was square root transformed to improve the stability of residual variance across fitted values. The model was specified as: where *Y_ijklm_* is lesion size at inoculation site *_m_* within pod of clone *i* inoculated with isolate *j* in bioassay *k*; μ is the overall intercept; *Ci* and *Ii* are the fixed effects of clone and isolate; and *CI_ij_* is their fixed interaction. The random intercept *Bk* accounts for variation among bioassays, while *P_l(ijk)_* accounts for variation among uniquely identified pods and dependence among sites within the same pod. The term Ɛ*_ijklm_* is the residual error associated with inoculation site *_m_* within pod *_l_* expressed on the square root scale.

The model was fitted by restricted maximum likelihood (REML). Fixed effects were evaluated using Type III analysis of variance with Satterthwaite degrees of freedom. The bioassay and pod random intercepts were evaluated using REML likelihood ratio tests, comparing the full model with reduced models that omitted each random intercept in turn while retaining the same fixed effects. These tests used approximate χ ^2^ reference distributions (Kuznetsova et al. 2017). The fitted model was checked for singularity. Residual versus fitted, scale location, and quantile–quantile plots were examined to assess model assumptions and compare residual behavior before and after transformation (Supplementary Fig. S2).

Estimated marginal means for clones and isolates were calculated with weighting proportional to the observed cell frequencies. Consequently, these summaries reflect the screening coverage of each clone or isolate rather than a common, equally represented panel. Estimates were back transformed to millimeters, and their covariance matrix was approximated on the response scale using the delta method. Confidence intervals and pairwise comparisons were then calculated on that scale. Pairwise comparisons among clones and among isolates were adjusted using Tukey’s method at α = 0.05 and summarized with compact letter displays. Shared letters indicate that a difference was not detected rather than equivalence between responses. Back-transformed estimates and their 95% confidence intervals are presented in Supplementary Tables S4 and S5; these estimates should not be interpreted as arithmetic means of the original observations. Model estimates for individual clone by isolate combinations were obtained separately and back transformed for presentation in the interaction plot.

Lesion expansion status was analyzed as a complementary binary response using sites with recorded reaction scores of 0 to 3 from the retained clones and isolates. Scores 0 and 1 were classified as no lesion expansion detected, whereas scores 2 and 3 were classified as lesion expansion observed. This classification was based on the recorded reaction phenotype rather than a lesion size threshold. The response was modeled using a binomial generalized linear mixed model with a logit link, coding lesion expansion observed as 1 and no lesion expansion detected as 0. Clone and isolate were included as fixed effects, with bioassay and the uniquely identified pod as random intercepts. A clone by isolate interaction was not included because outcome representation within combinations was sparse; the binary model therefore evaluated clone and isolate effects without estimating their interaction, and the interaction was assessed with the continuous lesion-size model.

The binary model was fitted using the bobyqa optimizer. Clone and isolate effects were evaluated using Type II Wald χ² tests. Estimated marginal probabilities of lesion expansion were obtained with 95% confidence intervals, and pairwise comparisons were performed on the logit scale with Tukey adjustment at α = 0.05. Model fit was examined using a binned residual plot and a Pearson overdispersion ratio.

Lesion size distributions and model estimates were displayed using boxplots, individual observations, and estimated marginal means. In these figures, a dashed reference line at 10 mm indicates the approximate width of the inoculation patch and was not used to classify lesion expansion or resistance. The clone by isolate interaction was illustrated using an interaction plot, and estimated probabilities of lesion expansion were presented in a combined supplementary figure. Package version information was recorded with each analysis run, and achieved P values are reported throughout.

## RESULTS

### Contrasting cacao responses to *Phytophthora* spp. inoculation

The inoculation assays revealed broad variation in lesion size among cacao clones in response to different *Phytophthora* species and isolates. We evaluated 25 cacao clones (Table 1) against 13 isolates representing *P. palmivora*, *P. megakarya*, and *P. theobromicola* (Table 2) across ten bioassays. Pod availability prevented a fully balanced screen, but 20 of the 25 clones were evaluated against more than six isolates, and observations were obtained for 241 of the 325 possible clone by isolate combinations (Supplementary Table S1). Across the full set of screened materials, lesion size at 4 days after inoculation ranged from 0 to 81.5 mm (Supplementary Fig. S1). Unadjusted mean lesion size among clones ranged from 10.34 mm for PENTAGONA-17 to 48.50 mm for PMCT-31, a 4.7-fold difference, with CCN-10 (39.67 mm) and NA-246 (39.22 mm) also near the upper end (Table 1). In this assay, isolate aggressiveness was assessed by mean lesion size at 4 days after inoculation, with larger lesions indicating greater aggressiveness. Across isolates, unadjusted mean lesion size ranged from 12.83 mm for *P. palmivora* Phy-VL to 43.87 mm for *P. megakarya* GH34, a 3.4-fold difference (Table 2). These unadjusted means reflect unequal clone by isolate coverage and therefore are not model-adjusted rankings.

Several patterns in the broader panel were biologically informative even when coverage was insufficient for model-based comparison. SCA-6 was included because it has been reported as a source of resistance. It had a small mean lesion size with *P. palmivora* GH49 (9.4 mm), but much larger means with *P. megakarya* ZTH0466 (42.4 mm) and GH34 (49.2 mm). Its mean with *P. palmivora* NPP3 was similarly large (48.4 mm), although this observation was based on one pod and is therefore exploratory. POUND-19-A was evaluated against seven isolates representing all three species. Four combinations met the minimum replication threshold, one fewer than required for the clone to enter the inferential analysis. It had the smallest observed clone mean with *P. megakarya* ZTH0466 (3.9 mm) and the second smallest mean among the 24 clones evaluated with *P. megakarya* GH34 (19.0 mm). Its mean with *P. palmivora* NPP3 was also small (6.3 mm), but this observation was based on one pod and is therefore exploratory. In contrast, PMCT-31 developed large lesions with each of the three isolates evaluated, one from each species, with means from 39.0 to 52.6 mm (Supplementary Table S1). These descriptive observations identify clone by isolate combinations that merit replicated evaluation.

### A continuum of lesion development among cacao clones

Lesion size differed among the 17 retained cacao clones and among the 12 *Phytophthora* isolates (Supplementary Table S1, bold entries in Panel A), and clone responses depended on the isolate: clone, F_(16, 449.2)_ = 25.16; isolate, F_(11, 416.0)_ = 32.90; and clone by isolate, F_(150, 465.4)_ = 1.72; all P < 0.001 (Table 3). Likelihood ratio tests supported retaining the pod random intercept, χ²(1) = 1,123.7, P < 0.001, and the bioassay random intercept, χ²(1) = 8.6, P = 0.003. Bioassay accounted for 3.7% of the random plus residual variance on the square-root scale, compared with 47.1% for the pod intercept and 49.2% for residual variation. Thus, variation among bioassays was detectable but comparatively small. Model-adjusted clone and isolate responses are presented in Figs. 1 and 2, respectively, and the clone by isolate interaction is illustrated in Fig. 3.

**Fig. 1.**
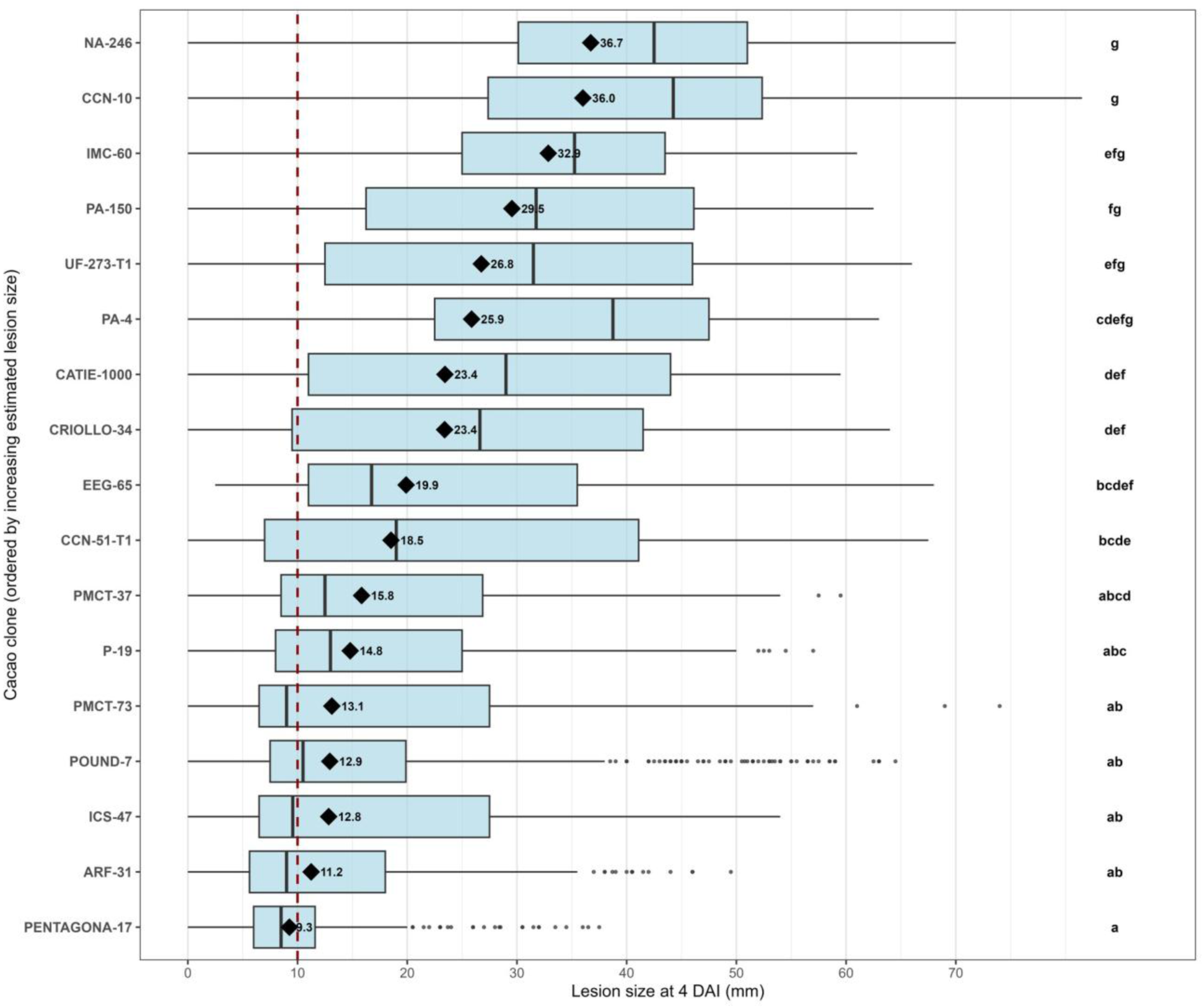
Estimated lesion size among cacao clones inoculated with *Phytophthora* spp. Boxplots show the distribution of site-level lesion size at 4 days after inoculation (DAI); diamonds mark the estimated marginal means (labeled, mm) from the lesion-size model, and lowercase letters denote Tukey-adjusted groups (α = 0.05); shared letters indicate that a difference was not detected. Clones are ordered by increasing estimated lesion size. The dashed line at 10 mm marks the approximate inoculation-patch width as a physical reference and was not used as a response-category threshold. The estimated marginal means with their 95% confidence intervals are tabulated in Supplementary Table S4.

**Fig. 2.**
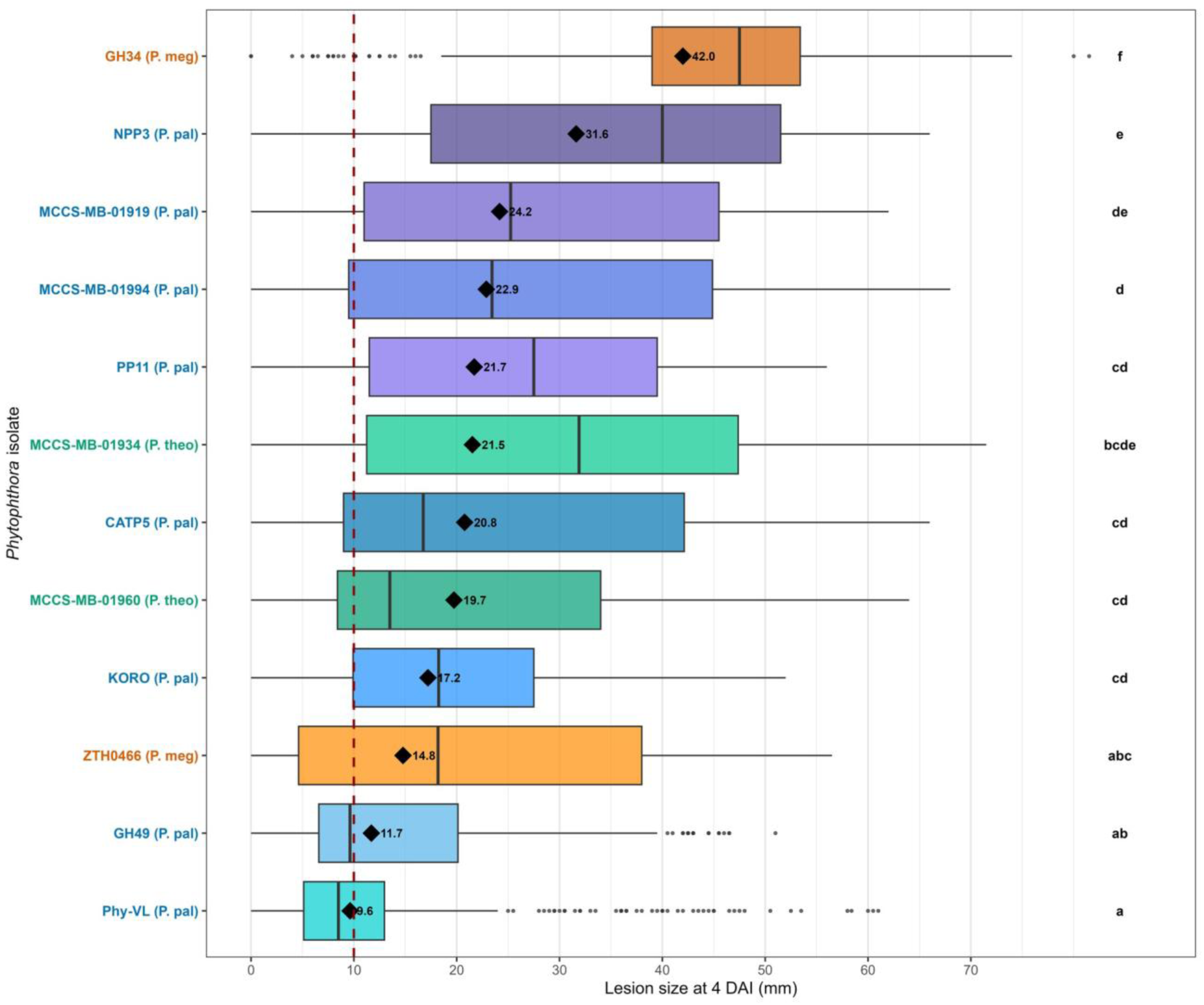
Aggressiveness of *Phytophthora* isolates on cacao pods, measured as estimated lesion size. Boxplots show site-level lesion size at 4 days after inoculation (DAI) across clones; diamonds mark the estimated marginal means (labeled, mm), and lowercase letters denote Tukey-adjusted groups (α = 0.05); shared letters indicate that a difference was not detected. Isolate labels are colored by species (*P. palmivora*, blue; *P. megakarya*, orange; *P. theobromicola*, green; P. pal, P. meg, and P. theo abbreviate the species in the axis labels). Isolates are ordered by increasing estimated lesion size. The dashed line at 10 mm marks the approximate inoculation-patch width as a physical reference and was not used as a response-category threshold. The estimated marginal means with their 95% confidence intervals are tabulated in Supplementary Table S5.

**Fig. 3.**
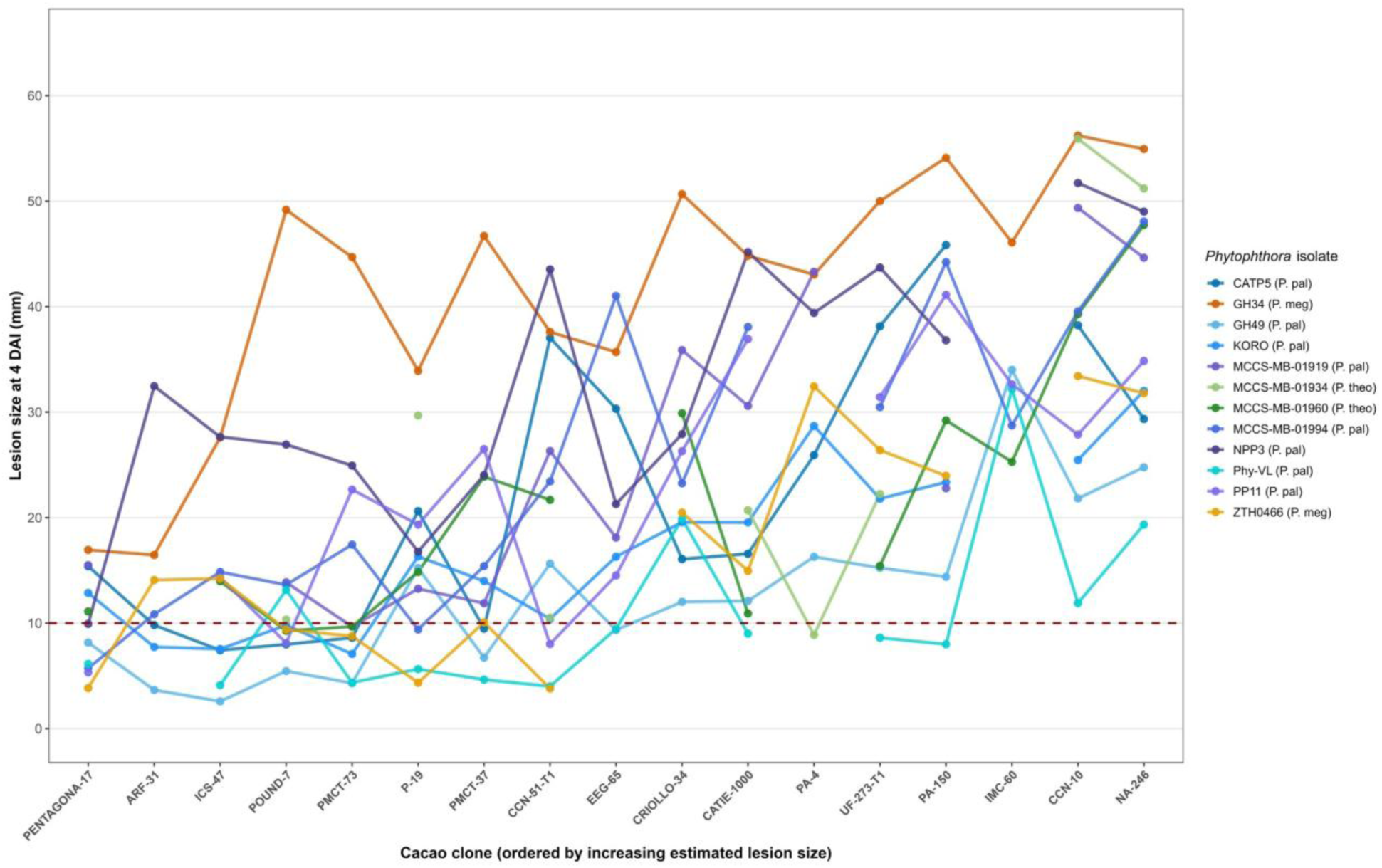
Interaction plot of cacao clone by *Phytophthora* isolate for lesion size. Each line connects the estimated marginal lesion size (mm) at 4 days after inoculation (DAI) of one *Phytophthora* isolate across the cacao clones; lines are colored by isolate and grouped by species (P. pal, *P. palmivora*; P. meg, *P. megakarya*; P. theo, *P. theobromicola*). Clones are ordered by increasing overall estimated lesion size. Lines display model estimates for the observed combinations; the overall clone by isolate interaction was evaluated using the fixed-effect test reported in Table 3. Gaps indicate clone by isolate combinations not represented in the retained dataset after filtering (Supplementary Table S1). The dashed line at 10 mm marks the approximate inoculation-patch width as a physical reference and was not used as a response-category threshold.

**Table 3.** Effects of cacao clone, *Phytophthora* isolate, and their interaction on lesion size (Type III analysis of variance).

| Source | Sum of squares | Mean square | Numerator df | Denominator df | *F* | *P* |
| --- | --- | --- | --- | --- | --- | --- |
| **Clone** | 472.4 | 29.53 | 16 | 449.2 | 25.16 | <0.001 |
| **Isolate** | 424.7 | 38.61 | 11 | 416.0 | 32.90 | <0.001 |
| **Clone × isolate** | 303.5 | 2.02 | 150 | 465.4 | 1.72 | <0.001 |

Linear mixed model of square-root-transformed lesion size at 4 days after inoculation, with cacao clone, *Phytophthora* isolate, and their interaction as fixed effects and bioassay and biological replicate pod as random intercepts; sums of squares are on the square-root scale. Denominator degrees of freedom (df) were approximated using Satterthwaite’s method. *P* values are achieved probabilities. × denotes the interaction term.

Estimated marginal means for lesion size ranged from 9.3 to 36.7 mm, an approximately fourfold difference (Fig. 1; Supplementary Table S4). PENTAGONA-17 had the smallest estimated lesion size (9.3 mm), followed by ARF-31 (11.2 mm), ICS-47 (12.8 mm), POUND-7 (12.9 mm), and PMCT-73 (13.1 mm). Intermediate estimates included P-19 (14.8 mm), PMCT-37 (15.8 mm), CCN-51-T1 (18.5 mm), and EEG-65 (19.9 mm). The largest estimated lesion sizes were found in NA-246 (36.7 mm), CCN-10 (36.0 mm), and IMC-60 (32.9 mm).

The five clones with the smallest estimated lesion sizes (PENTAGONA-17, ARF-31, ICS-47, POUND-7, and PMCT-73) differed significantly from the three with the largest (NA-246, CCN-10, and IMC-60), distinguishing the two ends of the distribution in Tukey-adjusted comparisons. Differences within either set were not detected (Fig. 1; Supplementary Table S4).

### Differences in aggressiveness among *Phytophthora* isolates

Aggressiveness varied widely among the 12 retained isolates. *P. palmivora* Phy-VL had the smallest estimated marginal lesion size (9.6 mm), and *P. megakarya* GH34 had the largest (42.0 mm), a 4.4-fold difference (Fig. 2; Supplementary Table S5). The contrast within *P. megakarya* was pronounced: its other isolate, ZTH0466, was near the lower end of the distribution at 14.8 mm. Among the eight *P. palmivora* isolates, Phy-VL and GH49 (11.7 mm) had the smallest estimates, NPP3 had the largest (31.6 mm), and the remaining five isolates had estimates between 17.2 and 24.2 mm. The two retained *P. theobromicola* isolates fell within this middle interval, with estimates of 19.7 mm for MCCS-MB-01960 and 21.5 mm for MCCS-MB-01934. These patterns describe variation among the isolates tested, whose species identities are given in Table 2, and do not constitute a formal comparison among species.

*P. megakarya* GH34 was the most aggressive isolate overall and differed significantly from every other isolate in Tukey-adjusted comparisons. Differences among *P. palmivora* Phy-VL, GH49, and *P. megakarya* ZTH0466 were not detected, and all three shared a compact letter. Both *P. theobromicola* isolates also shared letters with several *P. palmivora* isolates in the middle of the distribution. These marginal estimates summarize overall isolate aggressiveness across the observed clone and replication coverage, while differences among isolates depended on the clone (Table 3; Fig. 3).

### Clone responses depend on the isolate

Differences in lesion size among clones depended on the isolate, and differences among isolates depended on the clone (F_(150, 465.4)_ = 1.72, P < 0.001; Table 3). The fitted responses for the 178 retained combinations illustrate this interaction (Fig. 3), with gaps where combinations were not represented after filtering (Supplementary Table S1, Panel A). PENTAGONA-17 ranged from 3.8 mm with *P. megakarya* ZTH0466 to 16.9 mm with *P. megakarya* GH34 across 11 isolates. Other clones with lower or intermediate marginal estimates reached larger values with particular isolates, including PMCT-73 with *P. megakarya* GH34 (44.7 mm) and CCN-51-T1 with *P. palmivora* NPP3 (43.5 mm).

Relative clone ordering also changed among isolates. PENTAGONA-17 had a smaller estimate than POUND-7 with *P. megakarya* GH34 (16.9 versus 49.2 mm), but a larger estimate with *P. palmivora* CATP5 (15.4 versus 8.0 mm). Similarly, CCN-10 had a smaller estimate than NA-246 with *P. palmivora* MCCS-MB-01994 (39.5 versus 48.1 mm), but a larger estimate with *P. palmivora* CATP5 (38.2 versus 29.3 mm). These fitted point estimates illustrate the interaction descriptively; individual combinations or interaction departures were not tested separately. Separate views of the same fitted estimates for each species are provided in Supplementary Figs. S3 to S5.

### Reaction phenotypes and the likelihood of lesion expansion

All four reaction phenotypes were recorded among the scored inoculation sites. A typical expanding lesion (score 3) was the most frequent phenotype (50.6%), followed by localized darkening or necrosis confined to the inoculation area without lesion expansion (score 1; 25.8%), localized necrosis followed by an expanding lesion (score 2; 17.7%), and no visible reaction (score 0; 5.9%). For the complementary analysis, scores 0 and 1 represented no lesion expansion detected, and scores 2 and 3 represented lesion expansion observed. The observed proportion of sites with lesion expansion ranged from 38.2% for PENTAGONA-17 to 94.1% for NA-246 among clones and from 34.7% for *P. palmivora* Phy-VL to 96.6% for *P. megakarya* GH34 among isolates (Supplementary Tables S2 and S3). Within PENTAGONA-17, 89 sites showed localized necrosis without expansion, score 1, and 22 showed localized necrosis followed by expansion, score 2.

The probability of lesion expansion differed among cacao clones and among *Phytophthora* isolates in the binomial generalized linear mixed model (Type II Wald tests: clone, χ²(16) = 167.70, P < 0.001; isolate, χ²(11) = 161.68, P < 0.001).

PENTAGONA-17 had the lowest model-estimated probability of lesion expansion (0.296), whereas NA-246 had the highest (0.995). P-19 (0.740) and PMCT-37 (0.741) had intermediate estimates. PENTAGONA-17 had a significantly lower probability than the eight clones with estimates of 0.927 or higher, whereas NA-246 had a significantly higher probability than 11 of the other 16 clones. Many differences among clones with intermediate estimates were not detected in Tukey-adjusted comparisons (Supplementary Fig. S6A; Supplementary Table S2). A lack of detected differences does not imply equivalence.

Among isolates, *P. palmivora* Phy-VL had the lowest estimated probability of lesion expansion (0.207), whereas *P. megakarya* GH34 had the highest (0.997). Intermediate probabilities were estimated for *P. palmivora* CATP5 (0.688) and *P. megakarya* ZTH0466 (0.733). The two retained *P. theobromicola* isolates also had intermediate estimates: 0.729 for MCCS-MB-01934 and 0.823 for MCCS-MB-01960. Neither differed detectably from *P. palmivora* CATP5 or *P. megakarya* ZTH0466. *P. palmivora* Phy-VL had a significantly lower probability than every other isolate except *P. palmivora* GH49 and *P. theobromicola* MCCS-MB-01934, whereas *P. megakarya* GH34 had a significantly higher probability than every other isolate except *P. palmivora* MCCS-MB-01919 (Tukey-adjusted comparisons; Supplementary Fig. S6B; Supplementary Table S3). A clone by isolate interaction was not fitted for this response, so the binary model evaluated overall clone and isolate effects only. Lesion expansion status describes whether expansion was observed at a site, whereas the continuous analysis describes the lesion size attained at 4 days after inoculation.

## Discussion

This study evaluated the response of 25 cacao clones against 13 *Phytophthora* isolates representing three species, *P. palmivora*, *P. megakarya*, and *P. theobromicola*, under a common detached pod assay. Within the retained panel, lesion size differed among clones and isolates, and clone responses depended on the isolate (Table 3; Figs. 1 to 3). These findings connect the search for resistance in cacao with the diversity of the pathogens causing black pod disease. The broader panel also preserved preliminary observations that can guide the next stages of screening (Supplementary Table S1).

### Cacao responses and potential sources of resistance

Differences in lesion development among clones identified several candidates for black pod resistance breeding. PENTAGONA-17, ARF-31, ICS-47, POUND-7, and PMCT-73 had the five smallest overall estimated lesion sizes, whereas NA-246, CCN-10, and IMC-60 had the three largest (Fig. 1; Supplementary Table S4). PENTAGONA-17 stood out because its fitted lesion sizes remained between 3.8 and 16.9 mm across 11 isolates representing all three species (Fig. 3; Supplementary Table S1). It also had the lowest estimated probability of lesion expansion (Supplementary Fig. S6A; Supplementary Table S2). The other four candidates generally developed small lesions with most evaluated isolates, although some isolates produced larger lesions. Their representation in the fitted analysis differed: ARF-31 was represented by *P. palmivora* and *P. megakarya*, whereas ICS-47, POUND-7, and PMCT-73 were represented by all three species (Fig. 3; Supplementary Table S1).

Phillips et al. (2009) classified ARF-31, ICS-47, and POUND-7 as highly resistant to *P. palmivora* in CATIE, using artificial inoculations of detached pods. The results showed that these clones generally developed small lesions with most tested isolates (Fig. 3). These findings extend earlier evidence beyond *P. palmivora*.

Two of these candidates are also offspring of established resistance sources. ARF-31 is SCA-6 × CC-42, and PMCT-73 is POUND-7 × UF-667 (Table 1). POUND-7 has already been used as a black pod resistance parent in breeding, including the population studied by Gutiérrez et al. (2021). ARF-31 and PMCT-73 therefore represent descendants of established sources that developed small lesions with most isolates in our screen. Evaluating their progenies would help establish their value as parents for black pod resistance breeding.

The search for additional resistance sources can also draw on less used cacao germplasm. Upper Amazon germplasm has provided useful parents for black pod resistance (Nyassé et al. 2007), while numerous clones resistant to *P. palmivora* have been identified within the Guiana genetic group (Thévenin et al. 2012). Iwaro et al. (2006) further found a higher proportion of resistant accessions among wild than cultivated cacao in the collection they evaluated.

### Variation in aggressiveness among isolates

Lesion development revealed substantial variation in aggressiveness among isolates of the same species. The two *P. megakarya* isolates occupied opposite portions of the isolate ranking: GH34 had the largest estimated lesion size in the retained panel, at 42.0 mm, whereas ZTH0466 was among the three smallest, at 14.8 mm (Fig. 2; Supplementary Table S5). Differences among *P. megakarya* isolates have also been detected in leaf inoculation studies (Nyassé et al. 1995) and among selected multilocus genotypes evaluated by Djeumekop et al. (2026). Together, these results show that the response to one *P. megakarya* isolate does not describe the pathogen variation represented by other isolates of the same species.

The *P. palmivora* isolates also differed in aggressiveness. NPP3, had the largest estimated lesion size within this species, whereas Phy-VL and GH49 were at the lower end of the isolate ranking. NPP3 produced larger lesions than *P. megakarya* ZTH0466 but smaller lesions than GH34 (Fig. 2; Supplementary Table S5), showing that the isolate rankings overlapped across species in this panel. Rodríguez-Polanco et al. (2020) correspondingly, reported differences in lesion development among five *P. palmivora* isolates in cacao pod tests. The variation observed within both species supports including several isolates when determining how cacao candidates respond to the pathogen diversity represented in a screening collection.

The two *P. theobromicola* isolates retained in the fitted analysis had intermediate lesion estimates within the range occupied by several *P. palmivora* isolates (Fig. 2; Supplementary Table S5). Decloquement et al. (2021) in contrast, found that their tested *P. theobromicola* isolates generally produced faster lesion expansion than *P. palmivora* on four cacao clones. Their comparison used wounded pods and a different response measure, whereas the present study used unwounded pods and estimated lesion size. Our unfiltered screening dataset also included a third *P. theobromicola* isolate, MCCS-MB-01952, with observations across eight clones but only one qualifying clone by isolate combination (Table 2; Supplementary Table S1). Extending replicated coverage within this species would help determine how the candidate clones respond to additional *P. theobromicola* isolates.

### Clone responses depended on the isolate

Variation among isolates also changed the response obtained from individual cacao clones. The significant clone by isolate interaction means that differences among clones were not the same for every isolate (Table 3). The overall clone estimates identify candidates that generally developed small lesions, while the fitted combinations show the limits of those overall profiles. POUND-7 provides a clear example. Its fitted lesion sizes remained between 5.4 and 13.9 mm with ten isolates, but reached 26.9 mm with NPP3 and 49.2 mm with GH34. Its relative position also changed: POUND-7 developed smaller lesions than PENTAGONA-17 with CATP5, but much larger lesions than PENTAGONA-17 with GH34 (Fig. 3). These fitted combinations illustrate the interaction, while Table 3 provides its formal statistical support.

SCA-6 provides a complementary example involving an established resistance source. This clone has been used as a resistant control in leaf disc screening (Thévenin et al. 2012), but its observed mean lesion size in our unfiltered dataset was 9.4 mm with GH49 and increased to 42.4 and 49.2 mm with the two *P. megakarya* isolates, ZTH0466 and GH34, respectively. NPP3 also produced a mean lesion size of 48.4 mm, although that observation came from one pod (Supplementary Table S1). Ali et al. (2016) similarly, observed contrasting responses of attached SCA-6 pods to *P. palmivora* GH49 and *P. megakarya* GH34, with wounding changing the outcome. Fister et al. (2020) also found different responses between the two established sources: POUND-7 developed smaller lesions than SCA-6 in their detached-leaf assay with one *P. palmivora* isolate. Together, these comparisons show that an established resistance source can retain small-lesion responses to some pathogen challenges while developing much larger lesions with others.

Earlier studies have also produced different interaction patterns. Nyassé et al. (1995) detected a clone by species interaction between *P. palmivora* and *P. megakarya* in leaf assays but no clone by isolate interaction among three *P. megakarya* isolates. Surujdeo-Maharaj et al. (2001) observed a five- to sixfold range in aggressiveness among ten *P. palmivora* isolates but no genotype by isolate interaction. (Nyadanu et al. 2012b) also detected differences between *P. palmivora* and *P. megakarya* but no progeny by species interaction in leaf-disc and detached-pod assays.

In contrast, Barreto et al. (2015) detected a genotype by *Phytophthora* treatment interaction when 262 progeny were evaluated in leaf discs using one isolate each of *P. palmivora*, *P. capsici*, and *P. citrophthora*. That study also identified ten progeny classified as resistant to all three treatments, illustrating that a panel-level interaction can coexist with individual materials that retain low disease scores across all tested challenges. These studies differed in their cacao materials, pathogen panels, tissues, and assay designs. Together, they show that whether clone rankings remain similar across pathogen challenges must be evaluated for each screening panel.

In our study, the interaction does not diminish the value of clones that developed small lesions with most isolates. Instead, it identifies the particular combinations in which that general pattern changed. Evaluating candidates against several isolates is therefore important for identifying clones that generally develop small lesions while also revealing particular clone and isolate combinations that require further investigation.

### From localized reactions to the infection process

Lesion expansion status added a second view of the pod response. The probability of lesion expansion differed among clones and isolates, and the five clones with the smallest estimated lesion sizes also had the five lowest point estimates for expansion probability (Fig. 1; Supplementary Fig. S6A; Supplementary Tables S2 and S4). These related patterns did not make the two responses interchangeable. Lesion size described how extensively visible symptoms developed, whereas expansion status distinguished sites with an expanding lesion from sites where no expansion was detected under the scoring protocol. Localized necrosis was especially informative because it did not always have the same outcome. In PENTAGONA-17, for example, localized necrosis remained confined at some inoculation sites but was followed by lesion expansion at others. Because localized necrosis could remain confined or precede later lesion expansion, this phenotype raises questions about whether infection occurred and how pathogen development was related to the visible symptoms.

Iwaro et al. (1997b) distinguished lesion establishment from lesion development after penetration by measuring lesion frequency and lesion size separately. Ali et al. (2016) observed localized necrotic flecking on unwounded attached SCA-6 pods inoculated with *P. palmivora* and described it as resembling a hypersensitive response. Their observation provides a useful visual comparison with our localized phenotype, but appearance alone cannot identify the underlying process. We did not measure pathogen entry or colonization, and the score 2 phenotype shows that localized necrosis could also precede an expanding lesion. The present reactions therefore should not be interpreted as penetration resistance or assigned to a hypersensitive response.

Resolving these alternatives will require time course experiments combining microscopy or histology with measurements of pathogen biomass or viability and symptom development. A histological study associated variation in pod husk anatomy and lignification with contrasting black pod responses among cacao genotypes (Nyadanu et al. 2012a). Transcriptomic studies have compared early leaf responses in cacao genotypes differing in susceptibility to *P. megakarya* (Pokou et al. 2019) and *P. palmivora* (Winters et al. 2024). Expression of defense-associated genes has also been detected during susceptible infection (Ali et al. 2017), showing why host expression needs to be interpreted together with direct evidence of pathogen development. Applying these approaches to contrasting clone by isolate combinations could determine whether small lesion responses reflect unsuccessful infection, slower colonization, or restriction of pathogen growth after infection. Lesion expansion status therefore provides a useful phenotype for prioritizing those biological comparisons.

### Building the next stage of screening

The unfiltered screening dataset also retained preliminary leads that were excluded from the fitted panel because of limited coverage. POUND-19-A had an observed mean lesion size of 11.8 mm across seven isolates representing all three species, but only four clone by isolate combinations met the replication criterion. Its response to NPP3 came from one pod. POUND-19-A should therefore be prioritized for balanced follow-up rather than classified from the present observations. PNG-197 also had a small observed mean, 10.7 mm, but it was evaluated against only three *P. palmivora* isolates, with one pod per combination and all three pods evaluated in one bioassay (Table 1; Supplementary Table S1). These profiles identify materials for replicated screening, not resistance classifications.

Formal inference was based on 17 clones and 12 isolates represented in 178 of the 204 possible combinations, and some retained combinations were represented by only two independent pods. The retained isolates comprised eight *P. palmivora*, two *P. megakarya*, and two *P. theobromicola* isolates (Table 2; Supplementary Table S1). The six inoculation sites on each pod were subsamples of the same biological replicate rather than independent pods. The present design therefore supports comparisons among the sampled clones and isolates, but it was not designed to estimate species, genetic group, or geographic region effects. The lesion expansion model also estimated clone and isolate effects separately and did not test their interaction because combination level outcomes were too sparse (Supplementary Fig. S6; Supplementary Tables S2 and S3). Increasing the number of independent pods and balancing clone by isolate coverage would allow lesion size and lesion expansion to be evaluated together across a more comparable pathogen panel.

### From screening candidates to breeding and field evaluation

The candidates identified here were selected from their overall lesion estimates and their profiles across the sampled isolates (Figs. 1 and 3), but these responses do not by themselves establish their performance as parents. Nyadanu et al. (2012) found that parents with favorable general combining ability did not necessarily produce favorable specific crosses. Differences among progenies were also detected in detached-pod evaluations against *P. megakarya* by Ebaiarrey et al. (2024), while Ofori et al. (2023) found a positive association between detached-pod and field responses among 24 cacao families, together with parent- and cross-specific effects. Direct evaluation of progenies is therefore needed to determine the breeding value of ARF-31, PMCT-73, and the other candidates identified here.

Controlled pod assays can guide that process, but performance under field conditions must also be evaluated. Iwaro et al. (2005) found that detached pod rankings were associated with the average of three years of field observations, although the strength of the association varied among individual years. Efombagn et al. (2011) similarly found generally repeatable laboratory responses, but correspondence with field observations was not consistent across all comparisons. The candidates should therefore be tested in independent experiments with balanced clone by isolate coverage and evaluated across seasons under field conditions.

### Connecting resistance screening with pathogen surveillance

Expanding the cacao materials evaluated and maintaining an updated understanding of the pathogens causing black pod should proceed together. Molecular identification by Ali et al. (2016) distinguished black pod pathogens within collections from West Africa, while Decloquement et al. (2021) showed that isolates from diseased cacao pods in Brazil previously identified as *P. citrophthora* represented the distinct species *P. theobromicola*. Djeumekop et al. (2026) further combined repeated population sampling and genotyping of *P. megakarya* with leaf assays that detected differences among selected multilocus genotypes. These studies illustrate how species identification, population sampling, and phenotypic evaluation provide complementary information for selecting isolates for resistance screening. The collection locations recorded in Table 2 document the provenance of our isolate panel, but its uneven distribution among species and locations was not designed to test geographic effects. Collection location can therefore guide future sampling, whereas aggressiveness and clone responses must be measured experimentally.

This study links the search for resistance in cacao with variation in the pathogen challenge. PENTAGONA-17, ARF-31, ICS-47, POUND-7, and PMCT-73 generally developed small lesions across much of their evaluated isolate coverage, identifying candidates for continued resistance evaluation (Figs. 1 and 3; Supplementary Table S4). Aggressiveness differed among the isolates evaluated within species, particularly within *P. megakarya* and *P. palmivora* (Fig. 2; Supplementary Table S5). The significant clone by isolate interaction further showed that cacao responses to individual isolates differed among clones, explaining why a single isolate or one overall ranking cannot fully represent a candidate response (Table 3; Fig. 3). Lesion expansion status provided complementary observations that raise further questions about infection and colonization (Supplementary Fig. S6; Supplementary Tables S2 and S3). The next stage is to evaluate these candidates with more balanced isolate coverage and independent pods, followed by progeny and field evaluations. Connecting these experiments with continued pathogen sampling will allow resistance breeding to explore cacao diversity while remaining responsive to the diversity of the pathogens causing black pod disease.

## Supporting information

Sumplementary information

## Acknowledgments

The authors thank all the members of the Goss lab for their support during the inoculation process: Finnegan Corneliussen, Avril Rosano, and Sara M. Green. Special thanks to Robert Henfling from CATIE for his support with the cacao pods shipment from Costa Rica.

## LITERATURE CITED

Ali, S. S., Amoako-Attah, I., Bailey, R. A., Strem, M. D., Schmidt, M., Akrofi, A. Y., Surujdeo-Maharaj, S., Kolawole, O. O., Begoude, B. a. D., ten Hoopen, G. M., Goss, E., Phillips-Mora, W., Meinhardt, L. W., and Bailey, B. A. 2016. PCR-based identification of cacao black pod causal agents and identification of biological factors possibly contributing to Phytophthora megakarya’s field dominance in West Africa. Plant Pathol. 65:1095–1108. 10.1111/ppa.12496.

Ali, S. S., Shao, J., Lary, D. J., Strem, M. D., Meinhardt, L. W., and Bailey, B. A. 2017. Phytophthora megakarya and P. palmivora, Causal Agents of Black Pod Rot, Induce Similar Plant Defense Responses Late during Infection of Susceptible Cacao Pods. Front. Plant Sci. 8:169. 10.3389/fpls.2017.00169.

Barreto, M. A., Rosa, J. R. B. F., Holanda, I. S. A., Cardoso-Silva, C. B., Vildoso, C. I. A., Ahnert, D., Souza, M. M., Corrêa, R. X., Royaert, S., Marelli, J., Santos, E. S. L., Luz, E. D. M. N., Garcia, A. A. F., and Souza, A. P. 2018. QTL mapping and identification of corresponding genomic regions for black pod disease resistance to three Phytophthora species in Theobroma cacao L. Euphytica 214:188. 10.1007/s10681-018-2273-5.

Barreto, M. A., Santos, J. C. S., Corrêa, R. X., Luz, E. D. M. N., Marelli, J., and Souza, A. P. 2015. Detection of genetic resistance to cocoa black pod disease caused by three Phytophthora species. Euphytica 206:677–687. 10.1007/s10681-015-1490-4.

Baruah, I. K., Shao, J., Ali, S. S., Schmidt, M. E., Meinhardt, L. W., Bailey, B. A., and Cohen, S. P. 2024. Cacao pod transcriptome profiling of seven genotypes identifies features associated with post-penetration resistance to Phytophthora palmivora. Sci. Rep. 14:4175. 10.1038/s41598-024-54355-8.

Bates, D., Mächler, M., Bolker, B., and Walker, S. 2015. Fitting Linear Mixed-Effects Models Using lme4. J. Stat. Softw. 67:1–48. 10.18637/jss.v067.i01.

Boysen, O., Ferrari, E., Nechifor, V., and Tillie, P. 2023. Earn a living? What the Côte d’Ivoire–Ghana cocoa living income differential might deliver on its promise. Food Policy 114:102389. 10.1016/j.foodpol.2022.102389.

Brown, J. S., Phillips-Mora, W., Power, E. J., Krol, C., Cervantes-Martinez, C., Motamayor, J. C., and Schnell, R. J. 2007. Mapping QTLs for Resistance to Frosty Pod and Black Pod Diseases and Horticultural Traits in Theobroma cacao L. Crop Sci. 47:1851–1858. 10.2135/cropsci2006.11.0753.

Brugman, E., Wibowo, A., and Widiastuti, A. 2022. Phytophthora palmivora from Sulawesi and Java Islands, Indonesia, reveals high genotypic diversity and lack of population structure. Fungal Biol. 126:267–276. 10.1016/j.funbio.2022.02.004.

Clarke, E., Sherrill-Mix, S., and Dawson, C. 2023.ggbeeswarm: Categorical Scatter (Violin Point) Plots.

Dadzie, A. M., Obeng-Bio, E., Bukari, Y., Bediako, K. A., and Anyomi, E. W. 2026. Genetic variability and response of selected cocoa germplasm to black pod disease infection. Ecol. Genet. Genomics 40:100489. 10.1016/j.egg.2026.100489.

Daróczi, G., and Tsegelskyi, R. 2025. pander: An R “Pandoc” Writer.

Decloquement, J., Ramos-Sobrinho, R., Elias, S. G., Britto, D. S., Puig, A. S., Reis, A., da Silva, R. A. F., Honorato-Júnior, J., Luz, E. D. M. N., Pinho, D. B., and Marelli, J. P. 2021. Phytophthora theobromicola sp. nov.: A New Species Causing Black Pod Disease on Cacao in Brazil. Front. Microbiol. 12:537399. 10.3389/fmicb.2021.537399.

Djeumekop, M. M. N., Blondin, L., Herail, C., Ducamp, M., Vernière, C., ten Hoopen, G. M., and Neema, C. 2026. Genetic diversity of Phytophthora megakarya in relation to black pod disease spread in young cacao plantations in Cameroon. Eur. J. Plant Pathol. 10.1007/s10658-026-03287-2.

DuVal, A., Gezan, S. A., Mustiga, G., Stack, C., Marelli, J.-P., Chaparro, J., Livingstone, D., Royaert, S., and Motamayor, J. C. 2017. Genetic Parameters and the Impact of Off-Types for Theobroma cacao L. in a Breeding Program in Brazil. Front. Plant Sci. 8. 10.3389/fpls.2017.02059.

Ebaiarrey, H. E., Ngonkeu, E. L. M., Djoah, Y. T., and Efombagn, I. B. M. 2024. Assessing the Tolerance of Cocoa (Theobroma cacao L.) Progenies to the Black Pod Disease Caused by Phytophthora megakarya Bras. and Griff. J. Appl. Life Sci. Int. 27:13–28. 10.9734/jalsi/2024/v27i2638.

Efombagn, M. I. B., Bieysse, D., Nyassé, S., and Eskes, A. B. 2011. Selection for resistance to Phytophthora pod rot of cocoa ( Theobroma cacao L .) in Cameroon : Repeatability and reliability of screening tests and field observations. Crop Prot. 30:105–110. 10.1016/j.cropro.2010.10.012.

Ferguson, A. J., and Jeffers, S. N. 1999. Detecting Multiple Species of Phytophthora in Container Mixes from Ornamental Crop Nurseries. Plant Dis. 83:1129–1136. 10.1094/PDIS.1999.83.12.1129.

Fister, A. S., Leandro-Muñoz, M. E., Zhang, D., Marden, J. H., Tiffin, P., dePamphilis, C., Maximova, S., and Guiltinan, M. J. 2020. Widely distributed variation in tolerance to Phytophthora palmivora in four genetic groups of cacao. Tree Genet. Genomes 16:1. 10.1007/s11295-019-1396-8.

Fox, J., and Weisberg, S. 2019. An R Companion to Applied Regression. 3rd ed. Sage, Thousand Oaks, CA: Springer-Verlag New York.

García, J. F., Figueroa-Balderas, R., Puig, A. S., Kakati, I., Matson, M. E. H., Ali, S. S., Bailey, B. A., Marelli, J.-P., and Cantu, D. 2026. Intraspecies sequence-graph analysis of the Phytophthora theobromicola genome reveals a dynamic structure and variable effector repertoires. G3 GenesGenomesGenetics 16:jkaf256. 10.1093/g3journal/jkaf256.

Gitto, A. J., Schlathoelter, I., Herrera Corzo, M., Marelli, J. P., Goss, E. M., and Brawner, Jeremy T. 2026. Detached Pod-Zoospore Patch (DP-ZP) Phytophthora spp. Cacao Bioassay.

Graves, S., Piepho, H.-P., and Selzer, L. 2024. multcompView: Visualizations of Paired Comparisons.

Guest, D. 2007. Black Pod: Diverse Pathogens with a Global Impact on Cocoa Yield. Phytopathology® 97:1650–1653. 10.1094/PHYTO-97-12-1650.

Gutiérrez, O. A., Puig, A. S., Phillips-Mora, W., Bailey, B. A., Ali, S. S., Mockaitis, K., Schnell, R. J., Livingstone, D., Mustiga, G., Royaert, S., and Motamayor, J. C. 2021. SNP markers associated with resistance to frosty pod and black pod rot diseases in an F1 population of Theobroma cacao L. Tree Genet. Genomes 17:28. 10.1007/s11295-021-01507-w.

Hothorn, T., Bretz, F., and Westfall, P. 2008. Simultaneous Inference in General Parametric Models. Biom. J. 50:346–363. 10.1002/bimj.200810425.

Iwaro, A. D., Butler, D. R., and Eskes, A. B. 2006. Sources of Resistance to Phytophthora Pod Rot at the International CocoaGenebank, Trinidad. Genet. Resour. Crop Evol. 53:99–109. 10.1007/s10722-004-1411-1.

Iwaro, A. D., Sreenivasan, T. N., and Umaharan, P. 1997a. Foliar Resistance to Phytophthora palmivora as an Indicator of Pod Resistance in Theobroma cacao. Plant Dis. 81:619–624. 10.1094/PDIS.1997.81.6.619.

Iwaro, A. D., Sreenivasan, T. N., and Umaharan, P. 1997b. Phytophthora resistance in cacao (Theobroma cacao): Influence of pod morphological characteristics. Plant Pathol. 46:557–565. 10.1046/j.1365-3059.1997.d01-47.x.

Iwaro, A. D., Thévenin, J.-M., Butler, D. R., and Eskes, A. B. 2005. Usefulness of the Detached Pod Test for Assessment of Cacao Resistance to Phytophthora Pod Rot. Eur. J. Plant Pathol. 113:173–182. 10.1007/s10658-005-2929-6.

Kassambara, A. 2025. ggpubr: “ggplot2” Based Publication Ready Plots.

Kuznetsova, A., Brockhoff, P. B., and Christensen, R. H. B. 2017. lmerTest Package: Tests in Linear Mixed Effects Models. J. Stat. Softw. 82:1–26. 10.18637/jss.v082.i13.

Lenth, R. V., and Piaskowski, J. 2025. emmeans: Estimated marginal means, aka least-squares means. .

Marelli, J. P., Guest, D. I., Bailey, B. A., Evans, H. C., Brown, J. K., Junaid, M., Barreto, R. W., Lisboa, D. O., and Puig, A. S. 2019. Chocolate under threat from old and new cacao diseases. Phytopathology 109:1331–1343. 10.1094/PHYTO-12-18-0477-RVW.

Mata-Quirós, A., Arciniegas-Leal, A., Phillips-Mora, W., Meinhardt, L. W., and Zhang, D. 2017. Understanding the genetic structure and parentage of the clonal series of Cacao UF, CC, PMCT and ARF preserved in the International Cacao Collection at CATIE (IC3). In Lima, Peru.

Millard, S. P. 2013. EnvStats: An R Package for Environmental Statistics. New York: Springer. 10.1007/978-1-4614-8456-1.

Morales-Cruz, A., Ali, S. S., Minio, A., Figueroa-Balderas, R., García, J. F., Kasuga, T., Puig, A. S., Marelli, J.-P., Bailey, B. A., and Cantu, D. 2020. Independent Whole-Genome Duplications Define the Architecture of the Genomes of the Devastating West African Cacao Black Pod Pathogen *Phytophthora megakarya* and Its Close Relative *Phytophthora palmivora*. G3 GenesGenomesGenetics 10:2241–2255. 10.1534/g3.120.401014.

Motamayor, J. C., Lachenaud, P., da Silva e Mota, J. W., Loor, R., Kuhn, D. N., Brown, J. S., and Schnell, R. J. 2008. Geographic and Genetic Population Differentiation of the Amazonian Chocolate Tree (Theobroma cacao L). PLOS ONE 3:e3311. 10.1371/journal.pone.0003311.

Mucherino Muñoz, J. J., de Melo, C. A. F., Santana Silva, R. J., Luz, E. D. M. N., and Corrêa, R. X. 2021. Structural and Functional Genomics of the Resistance of Cacao to Phytophthora palmivora. Pathogens 10:961. 10.3390/pathogens10080961.

Nyadanu, D., Akromah, R., Adomako, B., Kwoseh, C., Lowor, S., Dzahini-Obiatey, H., Akrofi, A., Ansah, F. O., and Assuah, M. 2012a. Histological mechanisms of resistance to black pod disease in cacao (Theobroma cacao L.). J. Plant Sci. 7:39’54. 10.3923/jps.2012.39.54.

Nyadanu, D., Akromah, R., Adomako, B., Kwoseh, C., Lowor, S. T., Dzahini-Obiatey, H., Akrofi, A. Y., and Assuah, M. K. 2012b. Inheritance and general combining ability studies of detached pod, leaf disc and natural field resistance to Phytophthora palmivora and Phytophthora megakarya in cacao (Theobroma cacao L.). Euphytica 188:253–264. 10.1007/s10681-012-0717-x.

Nyassé, S., Cilas, C., Herail, C., and Blaha, G. 1995. Leaf inoculation as an early screening test for cocoa (*Theobroma cacao* L.) resistance to Phytophthora black pod disease. Crop Prot. 14:657–663. 10.1016/0261-2194(95)00054-2.

Nyassé, S., Efombagn, M. I. B., Kébé, B. I., Tahi, M., Despréaux, D., and Cilas, C. 2007. Integrated management of Phytophthora diseases on cocoa (Theobroma cacao L): Impact of plant breeding on pod rot incidence. Crop Prot. 26:40–45. 10.1016/j.cropro.2006.03.015.

Ofori, A., Padi, F. K., Amoako-Attah, I., Asare, E. K., Dadzie, A., and Bukari, Y. 2023. Genetic variation among cocoa (*Theobroma cacao* L.) families for resistance to black pod disease under field and laboratory conditions. Ecol. Genet. Genomics 28:100182. 10.1016/j.egg.2023.100182.

Pedersen, T. L. 2025. patchwork: The Composer of Plots.

Phillips, W., Castillo, J., Arciniegas, A., Mata, A., Sánchez, A., Leandro, M., Astorga, C., Motamayor, J., Guyton, B., Seguine, E., and others. 2009. Overcoming the main limiting factors of cacao production in Central America through the use of improved clones developed at CATIE. In Proceedings of the 16th International Cocoa Research Conference (Bali: COPAL Nigeria;) pp, , pp. 93–99.

Phillips-Mora, W., and Galindo, J. J. 1989. Método de inoculación y evaluación de la resistencia a Phytophthora palmivora en frutos de cacao (Theobroma cacao). Turrialba 39:488–496.

Pokou, D. N., Fister, A. S., Winters, N., Tahi, M., Klotioloma, C., Sebastian, A., Marden, J. H., Maximova, S. N., and Guiltinan, M. J. 2019. Resistant and susceptible cacao genotypes exhibit defense gene polymorphism and unique early responses to Phytophthora megakarya inoculation. Plant Mol. Biol. 99:499–516. 10.1007/s11103-019-00832-y.

Posit team. 2024. RStudio: Integrated Development Environment for R. . Pritchard, J. K., Stephens, M., and Donnelly, P. 2000. Inference of Population Structure Using Multilocus Genotype Data. Genetics 155:945–959.

Puig, A. S., Quintanilla, W., Matsumoto, T., Keith, L., Gutierrez, O. A., and Marelli, J.-P. 2021. Phytophthora palmivora Causing Disease on Theobroma cacao in Hawaii. Agriculture 11:396. 10.3390/agriculture11050396.

R Core Team. 2024.A language and environment for statistical computing. No Title. Revelle, W. 2025. psych: Procedures for psychological, psychometric, and personality research.

Risterucci, A. M., Paulin, D., Ducamp, M., N’Goran, J. A. K., and Lanaud, C. 2003. Identification of QTLs related to cocoa resistance to three species of Phytophthora. Theor. Appl. Genet. 108:168–174. 10.1007/s00122-003-1408-8.

Rodríguez Polanco, L., Carrero Gutiérrez, M. L., Parra, E. B., and Segura Amaya, J. D. 2022. Reaction of detached fruits from selected cocoa clones to artificial inoculation with Phytophthora palmivora. Acta Agronómica 71:186–194. 10.15446/acag.v71n2.88841.

Rodriguez-Medina, C., Arana, A. C., Sounigo, O., Argout, X., Alvarado, G. A., and Yockteng, R. 2019. Cacao breeding in Colombia, past, present and future. Breed. Sci. 69:373–382. 10.1270/jsbbs.19011.

Rodríguez-Polanco, E., Morales, J. G., Muñoz-Agudelo, M., Segura, J. D., and Carrero, M. L. 2020. Morphological, molecular and pathogenic characterization of Phytophthora palmivora isolates causing black pod rot of cacao in Colombia. Span. J. Agric. Res. 18:e1003. 10.5424/sjar/2020182-15147.

Schmidt, J. E., Puig, A. S., DuVal, A. E., and Pfeufer, E. E. 2023. Phyllosphere microbial diversity and specific taxa mediate within-cultivar resistance to Phytophthora palmivora in cacao. mSphere 8:e00013–23. 10.1128/msphere.00013-23.

Surujdeo-Maharaj, S., Umaharan, P., and Iwaro, A. D. 2001. A study of genotype-isolate interaction in cacao (Theobroma cacao L.): resistance of cacao genotypes to isolates of Phytophthora palmivora. Euphytica 118:295–303. 10.1023/A:1017516217662.

Tahi, G. M., Kébé, B. I., Sangare, A., Mondeil, F., Cilas, C., and Eskes, A. B. 2006. Foliar resistance of cacao (Theobroma cacao) to Phytophthora palmivora as an indicator of pod resistance in the field: Interaction of cacao genotype, leaf age and duration of incubation. Plant Pathol. 55:776–782. 10.1111/j.1365-3059.2006.01453.x.

Thévenin, J. M., Rossi, V., Ducamp, M., Doare, F., Condina, V., and Lachenaud, P. 2012. Numerous clones resistant to Phytophthora palmivora in the “Guiana” genetic group of Theobroma cacao L. PLoS ONE 7:1–6. 10.1371/journal.pone.0040915.

Wickham, H. 2025. forcats: Tools for Working with Categorical Variables (Factors). . Wickham, H. 2016. ggplot2: Elegant Graphics for Data Analysis. New York: Springer-Verlag.

Wickham, H., François, R., Henry, L., Müller, K., and Vaughan, D. 2023. dplyr: A Grammar of Data Manipulation. R package version 1.1.4. .

Winters, N. P., Wafula, E. K., Knollenberg, B. J., Hämälä, T., Timilsena, P. R., Perryman, M., Zhang, D., Sheaffer, L. L., Praul, C. A., Ralph, P. E., Prewitt, S., Leandro-Muñoz, M. E., Delgadillo-Duran, D. A., Altman, N. S., Tiffin, P., Maximova, S. N., dePamphilis, C. W., Marden, J. H., and Guiltinan, M. J. 2024. A combination of conserved and diverged responses underlies Theobroma cacao’s defense response to Phytophthora palmivora. BMC Biol. 22:38. 10.1186/s12915-024-01831-2.

Zhang, D., Martínez, W. J., Johnson, E. S., Somarriba, E., Phillips-Mora, W., Astorga, C., Mischke, S., and Meinhardt, L. W. 2012. Genetic diversity and spatial structure in a new distinct Theobroma cacao L. population in Bolivia. Genet. Resour. Crop Evol. 59:239–252. 10.1007/s10722-011-9680-y.

Zhang, D., and Motilal, L. 2016. Origin, Dispersal, and Current Global Distribution of Cacao Genetic Diversity. In Cacao Diseases, eds. Bryan A. Bailey and Lyndel W. Meinhardt. Cham, Switzerland: Springer International Publishing, pp. 3–31. 10.1007/978-3-319-24789-2_1.

