## Supplementary material for "Evaluating cacao pod resistance to black pod disease caused by three *Phytophthora* species": Sumplementary information

**Funding:** This research was funded by Mars Incorporated through a Sponsored Research Agreement with the University of Florida.

**The author(s) declare no conflict of interest**

Contents: Supplementary Tables S1 to S5 and Supplementary Figs. S1 to S6, cited in the main text. Supplementary Table S1 Panel A is presented in landscape orientation because of its 14 columns.

**Supplementary Table S1. Observed lesion size and biological replicate pod coverage for the complete cacao clone by *Phytophthora* isolate screening panel before inferential filtering. Panel A. Observed mean lesion size and biological replicate pod coverage by clone and isolate.**

| Cacao clone | Phy-VL | GH49 | KORO | MCCS-MB-01960 | ZTH0466 | CATP5 | PP11 | MCCS-MB-01994 | MCCS-MB-01919 | MCCS-MB-01934 | MCCS-MB-01952[a] | NPP3 | GH34 |
|---|---|---|---|---|---|---|---|---|---|---|---|---|---|
| **PENTAGONA-17** | **6.3 (5)** | **7.5 (3)** | **13.2 (4)** | **10.1 (2)** | **4.3 (4)** | **15.8 (2)** | **7.1 (2)** | **6.8 (4)** | **17.5 (3)** | – | – | **10.3 (3)** | **17.8 (4)** |
| **PNG-197[a]** | 9.8 (1) | 9.2 (1) | – | – | – | 12.9 (1) | – | – | – | – | – | – | – |
| **POUND-19-A[a]** | – | 8.5 (2) | – | 14.1 (2) | 3.9 (2) | 21.7 (1) | – | 10.8 (1) | – | – | – | 6.2 (1) | 19.0 (2) |
| **ARF-31** | – | **4.5 (4)** | **8.7 (2)** | 14.0 (1) | **16.9 (4)** | **10.5 (3)** | – | **10.3 (2)** | 13.9 (1) | 12.1 (1) | – | **35.6 (2)** | **21.3 (2)** |
| **PMCT-73** | **7.8 (3)** | **5.0 (4)** | **9.1 (3)** | **9.5 (5)** | **14.6 (3)** | **10.1 (4)** | **23.2 (4)** | **17.5 (3)** | **11.7 (3)** | – | – | **27.1 (3)** | **44.5 (4)** |
| **POUND-7** | **15.0 (5)** | **7.1 (7)** | **10.4 (6)** | **10.3 (6)** | **14.9 (6)** | **9.1 (6)** | **9.7 (6)** | **16.2 (6)** | **15.2 (6)** | **20.0 (2)** | 15.4 (1) | **31.2 (6)** | **51.1 (5)** |
| **HY-2714182[a]** | – | 1.3 (2) | – | 8.5 (1) | – | – | – | – | – | 11.5 (1) | – | – | 40.1 (2) |
| **P-19** | **7.7 (5)** | **15.6 (7)** | **16.9 (6)** | **15.2 (8)** | **5.1 (5)** | **23.2 (6)** | **21.2 (5)** | **15.0 (5)** | **14.1 (4)** | **34.0 (2)** | – | **19.2 (5)** | **34.2 (4)** |
| **ICS-47** | **5.6 (2)** | **4.5 (3)** | **9.8 (2)** | **13.4 (4)** | **17.7 (3)** | **8.6 (2)** | **21.3 (2)** | **17.2 (3)** | – | 50.5 (1) | 16.0 (1) | **31.6 (3)** | **32.9 (3)** |
| **PMCT-37** | **7.0 (4)** | **6.7 (3)** | **15.2 (4)** | **23.6 (4)** | **16.0 (3)** | **9.6 (3)** | **27.2 (4)** | **14.3 (3)** | **11.3 (4)** | – | 42.1 (1) | **25.6 (3)** | **45.0 (3)** |
| **COCA-3370-5[a]** | 9.6 (1) | 9.8 (1) | 17.8 (1) | – | 21.3 (1) | – | – | – | 17.2 (2) | 8.3 (1) | 21.8 (1) | 22.5 (2) | 43.6 (1) |
| **ARF-10[a]** | – | 7.9 (1) | – | 9.2 (1) | – | – | – | 31.2 (1) | – | – | – | – | 44.2 (1) |
| **CCN-51-T1** | **4.8 (4)** | **17.2 (4)** | **11.6 (4)** | **23.5 (5)** | **11.7 (3)** | **42.4 (4)** | **12.2 (3)** | **26.6 (4)** | **31.6 (4)** | **12.2 (2)** | – | **46.1 (3)** | **38.7 (4)** |
| **EEG-65** | **10.9 (3)** | **10.8 (3)** | **18.8 (3)** | 24.1 (1) | 29.8 (1) | **34.4 (3)** | **17.2 (3)** | **44.2 (2)** | **20.8 (2)** | – | 59.7 (2) | **26.5 (3)** | **38.7 (3)** |
| **CRIOLLO-34** | **22.0 (4)** | **13.4 (6)** | **19.5 (4)** | **30.2 (4)** | **25.9 (3)** | **17.6 (4)** | **31.4 (4)** | **28.7 (4)** | **37.8 (3)** | – | 9.8 (1) | **36.0 (4)** | **51.2 (3)** |
| **LCTEEN-37-F[a]** | 29.4 (1) | 15.2 (1) | 19.7 (1) | – | – | 48.2 (1) | 29.8 (1) | 11.1 (1) | – | – | – | 28.8 (1) | 38.0 (1) |
| **SCA-6[a]** | – | 9.4 (2) | 14.4 (1) | 9.4 (2) | 42.4 (2) | 20.2 (1) | – | – | 42.2 (1) | 21.0 (1) | – | 48.4 (1) | 49.2 (2) |
| **CATIE-1000** | **9.8 (3)** | **15.9 (4)** | **24.2 (3)** | **11.5 (3)** | **22.0 (5)** | **21.1 (4)** | **39.8 (4)** | **39.1 (4)** | **35.2 (3)** | **27.7 (2)** | – | **48.1 (4)** | **46.2 (3)** |
| **UF-273-T1** | **8.7 (3)** | **18.6 (4)** | **22.9 (3)** | **15.9 (4)** | **31.0 (2)** | **41.0 (4)** | **36.0 (2)** | **29.3 (3)** | 27.1 (1) | **29.1 (2)** | 19.7 (1) | **43.7 (4)** | **50.3 (4)** |
| **PA-150** | **9.0 (3)** | **14.7 (3)** | **23.1 (4)** | **30.1 (4)** | **23.9 (3)** | **44.8 (3)** | **42.1 (3)** | **44.2 (3)** | **27.8 (2)** | – | – | **36.8 (3)** | **53.1 (3)** |
| **IMC-60** | **35.9 (2)** | **32.5 (2)** | – | **25.5 (2)** | – | – | **34.8 (2)** | **27.5 (2)** | – | – | – | – | **43.6 (2)** |
| **PA-4** | 17.3 (1) | **21.4 (3)** | **33.3 (3)** | – | **40.5 (4)** | **33.7 (2)** | 35.8 (1) | – | **47.6 (2)** | **18.6 (4)** | – | **44.5 (2)** | **50.1 (2)** |
| **NA-246** | **22.3 (6)** | **25.6 (5)** | **32.9 (6)** | **45.5 (3)** | **35.8 (5)** | **31.3 (6)** | **38.5 (4)** | **47.9 (5)** | **47.0 (4)** | **53.8 (2)** | – | **49.9 (5)** | **55.8 (6)** |
| **CCN-10** | **18.2 (3)** | **25.0 (5)** | **27.1 (6)** | **39.4 (6)** | **36.8 (4)** | **41.2 (4)** | **32.9 (4)** | **44.1 (4)** | **47.8 (5)** | **59.0 (2)** | 56.5 (1) | **51.1 (4)** | **57.2 (5)** |
| **PMCT-31[a]** | – | 39.0 (3) | – | – | – | – | – | – | – | 52.6 (3) | – | – | 52.5 (4) |

(continued in Panels B and C)

### Supplementary Table S1 (continued). Panel B. Cacao clones (n = 25).

| Cacao clone | Mean (mm) | SD (mm) | Range (mm) | Site records (n) | Isolates evaluated (n) | Biological replicate pods represented (n) |
|---|---|---|---|---|---|---|
| **PENTAGONA-17** | 10.34 | 7.74 | 0.0–37.5 | 216 | 11 | 36 |
| **PNG-197[a]** | 10.67 | 2.46 | 5.5–13.5 | 18 | 3 | 3 |
| **POUND-19-A[a]** | 11.78 | 10.94 | 0.0–55.0 | 66 | 7 | 11 |
| **ARF-31** | 14.03 | 12.31 | 0.0–49.5 | 132 | 10 | 22 |
| **PMCT-73** | 16.46 | 15.40 | 0.0–74.0 | 234 | 11 | 39 |
| **POUND-7** | 16.74 | 16.23 | 0.0–64.5 | 408 | 13 | 68 |
| **HY-2714182[a]** | 17.16 | 20.10 | 0.0–56.0 | 36 | 4 | 6 |
| **P-19** | 17.36 | 12.95 | 0.0–57.0 | 366 | 12 | 62 |
| **ICS-47** | 18.02 | 16.12 | 0.0–57.5 | 174 | 12 | 29 |
| **PMCT-37** | 18.83 | 14.44 | 0.0–59.5 | 234 | 12 | 39 |
| **COCA-3370-5[a]** | 19.22 | 12.36 | 4.0–48.5 | 66 | 9 | 11 |
| **ARF-10[a]** | 23.12 | 18.26 | 0.0–49.0 | 24 | 4 | 4 |
| **CCN-51-T1** | 23.71 | 19.36 | 0.0–67.5 | 264 | 12 | 44 |
| **EEG-65** | 26.74 | 17.56 | 2.5–70.5 | 174 | 12 | 29 |
| **CRIOLLO-34** | 26.75 | 17.59 | 0.0–64.0 | 264 | 12 | 44 |
| **LCTEEN-37-F[a]** | 27.51 | 13.54 | 5.0–52.5 | 48 | 8 | 8 |
| **SCA-6[a]** | 28.23 | 18.11 | 0.0–52.5 | 78 | 9 | 13 |
| **CATIE-1000** | 28.61 | 17.47 | 0.0–59.5 | 252 | 12 | 42 |
| **UF-273-T1** | 29.73 | 17.03 | 0.0–66.0 | 222 | 13 | 37 |
| **PA-150** | 31.59 | 17.22 | 0.0–62.5 | 204 | 11 | 34 |
| **IMC-60** | 33.30 | 15.09 | 0.0–61.0 | 72 | 6 | 12 |
| **PA-4** | 33.54 | 17.24 | 0.0–63.0 | 144 | 10 | 24 |
| **NA-246** | 39.22 | 14.76 | 0.0–70.0 | 342 | 12 | 57 |
| **CCN-10** | 39.67 | 17.95 | 0.0–81.5 | 318 | 13 | 53 |
| **PMCT-31[a]** | 48.50 | 9.32 | 0.0–57.5 | 60 | 3 | 10 |

(continued in Panel C)

### Supplementary Table S1 (continued). Panel C. *Phytophthora* isolates (n = 13).

| Isolate | Mean (mm) | SD (mm) | Range (mm) | Site records (n) | Clones evaluated (n) | Biological replicate pods represented (n) |
|---|---|---|---|---|---|---|
| **Phy-VL** | 12.83 | 13.18 | 0.0–63.0 | 354 | 19 | 59 |
| **GH49** | 14.42 | 12.26 | 0.0–51.0 | 498 | 25 | 83 |
| **KORO** | 19.27 | 10.95 | 0.0–52.0 | 396 | 19 | 66 |
| **MCCS-MB-01960** | 20.33 | 15.30 | 0.0–64.0 | 408 | 20 | 68 |
| **ZTH0466** | 21.16 | 17.70 | 0.0–56.5 | 378 | 19 | 63 |
| **CATP5** | 24.88 | 18.02 | 0.0–66.0 | 384 | 20 | 64 |
| **PP11** | 26.21 | 15.44 | 0.0–56.0 | 323 | 17 | 54 |
| **MCCS-MB-01994** | 26.30 | 19.26 | 0.0–68.0 | 360 | 19 | 60 |
| **MCCS-MB-01919** | 27.53 | 17.57 | 0.0–62.0 | 295 | 17 | 50 |
| **MCCS-MB-01934** | 31.05 | 20.51 | 0.0–71.5 | 156 | 14 | 26 |
| **MCCS-MB-01952[a]** | 33.40 | 21.66 | 5.0–70.5 | 54 | 8 | 9 |
| **NPP3** | 34.66 | 18.37 | 0.0–66.0 | 372 | 20 | 62 |
| **GH34** | 43.87 | 14.64 | 0.0–81.5 | 438 | 24 | 73 |

Panel A cells report the arithmetic mean of the available site-level lesion-size values at 4 days after inoculation, in millimeters, followed in parentheses by the number of biological replicate pods represented; an en dash indicates that the combination was not evaluated; nc indicates a combination whose records lacked a calculable lesion-size value. Panels B and C report, for each clone and each isolate, the mean, standard deviation (SD), and range of the available site-level values, the number of site records, the number of partner levels evaluated, and the number of biological replicate pods represented. All values are descriptive summaries of the complete post-processing dataset (4,416 site records; 737 biological replicate pods) after pod processing and within-pod imputation and before inferential filtering; they are not model-adjusted estimates and should not be interpreted as rankings. Seven records lacked a calculable lesion-size value and were omitted from means, SD, and ranges. Values based on one pod are exploratory observations. Values in bold in Panel A identify the 178 clone by isolate combinations retained for the primary lesion-size model (17 clones, 12 isolates; 178 of the 204 possible combinations among them); every retained combination was represented by at least two pods, and these entries total 654 pods. In each of two retained combinations, P-19 × MCCS-MB-01919 and PMCT-37 × KORO, one pod lacked a calculable lesion-size response, so 652 pod groups contributed observations to the fitted model. [a] Level excluded from the inferential analyses because fewer than five clone by isolate combinations qualified (at least 12 site records after imputation, equivalent to at least two replicate pods). Supplementary Tables S4 and S5 give the model-adjusted estimates for the retained levels. Clones and isolates are ordered by increasing observed mean lesion size.

**Supplementary Table S2. Observed proportion and model estimated probability of lesion expansion by cacao clone.**

| Cacao clone | Scored sites (n) | No lesion expansion detected (n) | Lesion expansion observed (n) | Lesion expansion observed (%) | Estimated probability (95% CI) | Group |
|---|---|---|---|---|---|---|
| **PENTAGONA-17** | 165 | 102 | 63 | 38.2 | 0.296 (0.132–0.538) | a |
| **ARF-31** | 93 | 56 | 37 | 39.8 | 0.336 (0.121–0.649) | ab |
| **PMCT-73** | 227 | 118 | 109 | 48.0 | 0.485 (0.271–0.704) | ab |
| **POUND-7** | 391 | 201 | 190 | 48.6 | 0.485 (0.312–0.663) | ab |
| **ICS-47** | 156 | 74 | 82 | 52.6 | 0.511 (0.257–0.759) | ab |
| **P-19** | 337 | 140 | 197 | 58.5 | 0.740 (0.565–0.862) | abc |
| **PMCT-37** | 222 | 89 | 133 | 59.9 | 0.741 (0.521–0.883) | abc |
| **CCN-51-T1** | 247 | 92 | 155 | 62.8 | 0.772 (0.577–0.894) | abcd |
| **EEG-65** | 145 | 31 | 114 | 78.6 | 0.863 (0.629–0.959) | abcde |
| **PA-4** | 132 | 20 | 112 | 84.8 | 0.927 (0.758–0.981) | bcdef |
| **CRIOLLO-34** | 245 | 64 | 181 | 73.9 | 0.928 (0.834–0.971) | cde |
| **CATIE-1000** | 247 | 60 | 187 | 75.7 | 0.932 (0.837–0.973) | cde |
| **UF-273-T1** | 184 | 31 | 153 | 83.2 | 0.960 (0.889–0.986) | cdef |
| **PA-150** | 196 | 32 | 164 | 83.7 | 0.973 (0.921–0.991) | def |
| **CCN-10** | 304 | 33 | 271 | 89.1 | 0.976 (0.938–0.991) | ef |
| **IMC-60** | 70 | 9 | 61 | 87.1 | 0.981 (0.889–0.997) | cdef |
| **NA-246** | 339 | 20 | 319 | 94.1 | 0.995 (0.986–0.999) | f |

Lesion expansion observed corresponds to reaction scores 2 and 3, whereas no lesion expansion detected corresponds to scores 0 and 1. Columns provide the number of scored inoculation sites, counts for both binary categories, the observed percentage with lesion expansion, and the probability of lesion expansion estimated by the binomial generalized linear mixed model with its 95% confidence interval (CI). Confidence intervals were back transformed from the logit scale. Pairwise comparisons among clones were adjusted using Tukey's method on the logit scale at $\alpha = 0.05$. Shared letters indicate that a difference was not detected and do not demonstrate equivalence. Clones are ordered by increasing estimated probability.

**Supplementary Table S3. Observed proportion and model estimated probability of lesion expansion by *Phytophthora* isolate.**

| Isolate | Species | Scored sites (n) | No lesion expansion detected (n) | Lesion expansion observed (n) | Lesion expansion observed (%) | Estimated probability (95% CI) | Group |
|---|---|---|---|---|---|---|---|
| **Phy-VL** | *P. palmivora* | 303 | 198 | 105 | 34.7 | 0.207 (0.104–0.370) | a |
| **GH49** | *P. palmivora* | 400 | 205 | 195 | 48.8 | 0.433 (0.269–0.614) | ab |
| **CATP5** | *P. palmivora* | 342 | 133 | 209 | 61.1 | 0.688 (0.499–0.830) | bc |
| **MCCS-MB-01934** | *P. theobromicola* | 108 | 26 | 82 | 75.9 | 0.729 (0.406–0.914) | abcd |
| **ZTH0466** | *P. megakarya* | 321 | 136 | 185 | 57.6 | 0.733 (0.549–0.861) | bc |
| **MCCS-MB-01960** | *P. theobromicola* | 331 | 115 | 216 | 65.3 | 0.823 (0.657–0.918) | cd |
| **KORO** | *P. palmivora* | 355 | 103 | 252 | 71.0 | 0.883 (0.767–0.945) | cd |
| **MCCS-MB-01994** | *P. palmivora* | 320 | 98 | 222 | 69.4 | 0.900 (0.791–0.955) | cd |
| **PP11** | *P. palmivora* | 295 | 58 | 237 | 80.3 | 0.918 (0.819–0.965) | cd |
| **NPP3** | *P. palmivora* | 325 | 57 | 268 | 82.5 | 0.953 (0.895–0.980) | d |
| **MCCS-MB-01919** | *P. palmivora* | 250 | 31 | 219 | 87.6 | 0.974 (0.925–0.991) | de |
| **GH34** | *P. megakarya* | 350 | 12 | 338 | 96.6 | 0.997 (0.991–0.999) | e |

Lesion expansion observed corresponds to reaction scores 2 and 3, whereas no lesion expansion detected corresponds to scores 0 and 1. Columns provide the number of scored inoculation sites, counts for both binary categories, the observed percentage with lesion expansion, and the probability of lesion expansion estimated by the binomial generalized linear mixed model with its 95% confidence interval (CI). Confidence intervals were back transformed from the logit scale. Pairwise comparisons among isolates were adjusted using Tukey's method on the logit scale at $\alpha = 0.05$. Shared letters indicate that a difference was not detected and do not demonstrate equivalence. Isolates are ordered by increasing estimated probability.

### Supplementary Table S4. Model estimated marginal lesion size for each cacao clone.

| Cacao clone | Estimated lesion size (mm) | 95% CI (mm) | Group |
|---|---|---|---|
| **PENTAGONA-17** | 9.3 | 6.5–12.0 | a |
| **ARF-31** | 11.2 | 7.4–15.1 | ab |
| **ICS-47** | 12.8 | 9.3–16.4 | ab |
| **POUND-7** | 12.9 | 10.4–15.4 | ab |
| **PMCT-73** | 13.1 | 10.0–16.2 | ab |
| **P-19** | 14.8 | 12.0–17.6 | abc |
| **PMCT-37** | 15.8 | 12.4–19.3 | abcd |
| **CCN-51-T1** | 18.5 | 15.0–22.1 | bcde |
| **EEG-65** | 19.9 | 15.1–24.7 | bcdef |
| **CRIOLLO-34** | 23.4 | 19.5–27.4 | def |
| **CATIE-1000** | 23.4 | 19.3–27.6 | def |
| **PA-4** | 25.9 | 19.8–31.9 | cdefg |
| **UF-273-T1** | 26.8 | 22.2–31.3 | efg |
| **PA-150** | 29.5 | 24.7–34.4 | fg |
| **IMC-60** | 32.9 | 24.7–41.0 | efg |
| **CCN-10** | 36.0 | 31.3–40.7 | g |
| **NA-246** | 36.7 | 32.1–41.3 | g |

Estimated marginal means (EMM) of lesion size at 4 days after inoculation from the linear mixed model, calculated with weighting proportional to the observed cell frequencies; because coverage was incomplete, each estimate summarizes a clone over the clone by isolate combinations actually represented for it (Supplementary Table S1) rather than a common, equally represented panel. Means were back transformed to millimeters, and their covariance matrix was approximated on the response scale using the delta method; the 95% confidence intervals (CI) were calculated on that scale. Pairwise comparisons were adjusted using Tukey's method at $\alpha = 0.05$ and are summarized as compact-letter groups; shared letters indicate that a difference was not detected and do not demonstrate equivalence. These are the estimates plotted as diamonds in Fig. 1. Clones are ordered by increasing estimated lesion size.

**Supplementary Table S5. Model estimated marginal lesion size (aggressiveness) for each *Phytophthora* isolate.**

| Isolate | Species | Estimated lesion size (mm) | 95% CI (mm) | Group |
|---|---|---|---|---|
| **Phy-VL** | *P. palmivora* | 9.6 | 7.3–12.0 | a |
| **GH49** | *P. palmivora* | 11.7 | 9.4–14.0 | ab |
| **ZTH0466** | *P. megakarya* | 14.8 | 11.9–17.7 | abc |
| **KORO** | *P. palmivora* | 17.2 | 14.2–20.2 | cd |
| **MCCS-MB-01960** | *P. theobromicola* | 19.7 | 15.9–23.6 | cd |
| **CATP5** | *P. palmivora* | 20.8 | 17.4–24.1 | cd |
| **MCCS-MB-01934** | *P. theobromicola* | 21.5 | 15.9–27.2 | bcde |
| **PP11** | *P. palmivora* | 21.7 | 18.0–25.4 | cd |
| **MCCS-MB-01994** | *P. palmivora* | 22.9 | 19.2–26.5 | d |
| **MCCS-MB-01919** | *P. palmivora* | 24.2 | 20.1–28.2 | de |
| **NPP3** | *P. palmivora* | 31.6 | 27.5–35.8 | e |
| **GH34** | *P. megakarya* | 42.0 | 37.3–46.7 | f |

Computed exactly as described for Supplementary Table S4, for isolates: estimated marginal means (EMM) with 95% confidence intervals (CI) on the response scale and Tukey-adjusted compact-letter groups ($\alpha = 0.05$); shared letters indicate that a difference was not detected and do not demonstrate equivalence. These are the estimates plotted as diamonds in Fig. 2. Isolates are ordered by increasing estimated lesion size.

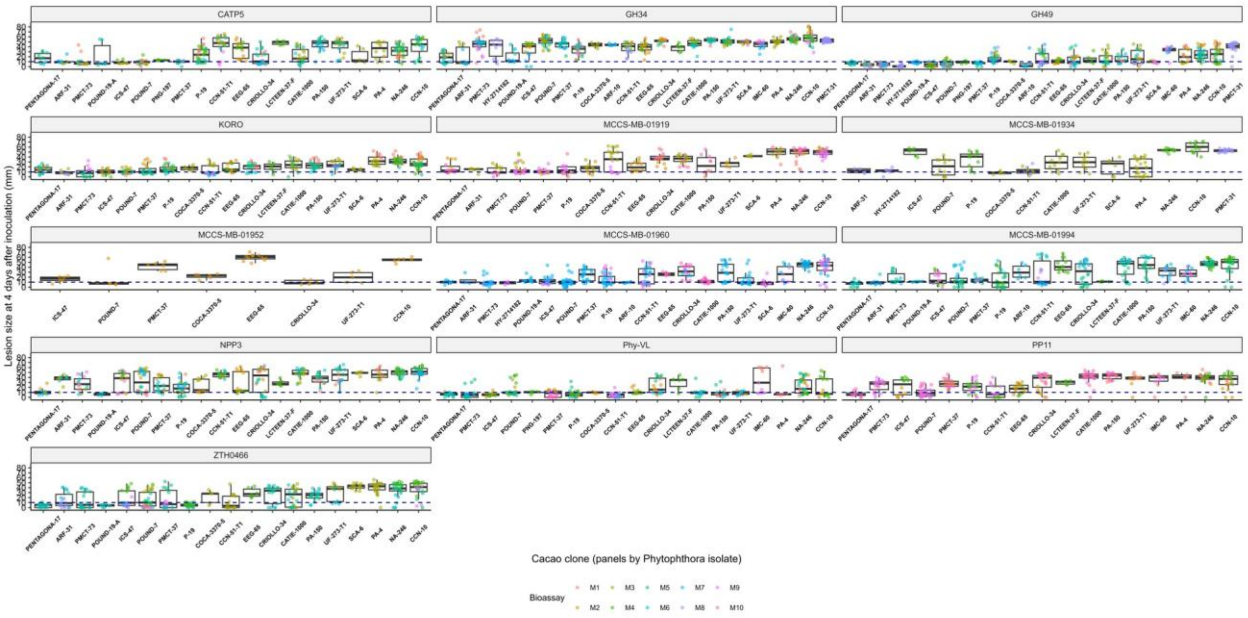


**Supplementary Fig. S1.** Distribution of site-level lesion size at 4 days after inoculation for all 25 cacao clones and 13 *Phytophthora* isolates in the processed dataset before imputation and before inferential filtering (4,416 site records; 265 records without a measurable lesion value are not shown). Each panel shows one isolate; boxplots summarize the observed values for each clone, and points show individual inoculation sites colored by bioassay (M1 to M10). The dashed line at 10 mm marks the approximate inoculation-patch width as a physical reference.

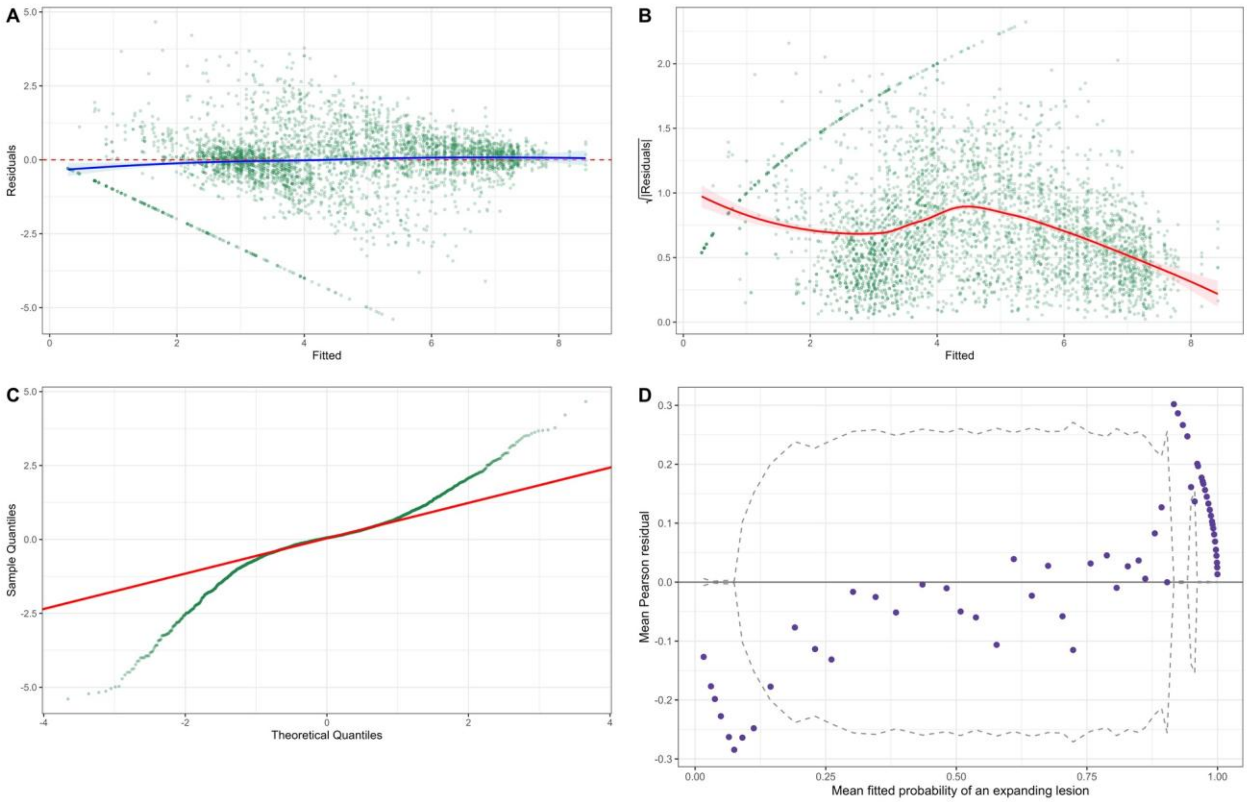


**Supplementary Fig. S2.** Residual diagnostics for the lesion-size model and the binomial lesion-expansion model. (A) Residuals versus fitted values, (B) scale–location plot (square root of absolute residuals versus fitted values), and (C) normal quantile–quantile plot for the linear mixed model of square-root-transformed lesion size; fitted values and residuals are on the square-root scale. (D) Binned residual plot for the binomial generalized linear mixed model of lesion expansion status: mean Pearson residual against mean fitted probability in 60 bins; dashed lines mark approximate ±2 standard error bounds; the Pearson dispersion ratio was 0.45.

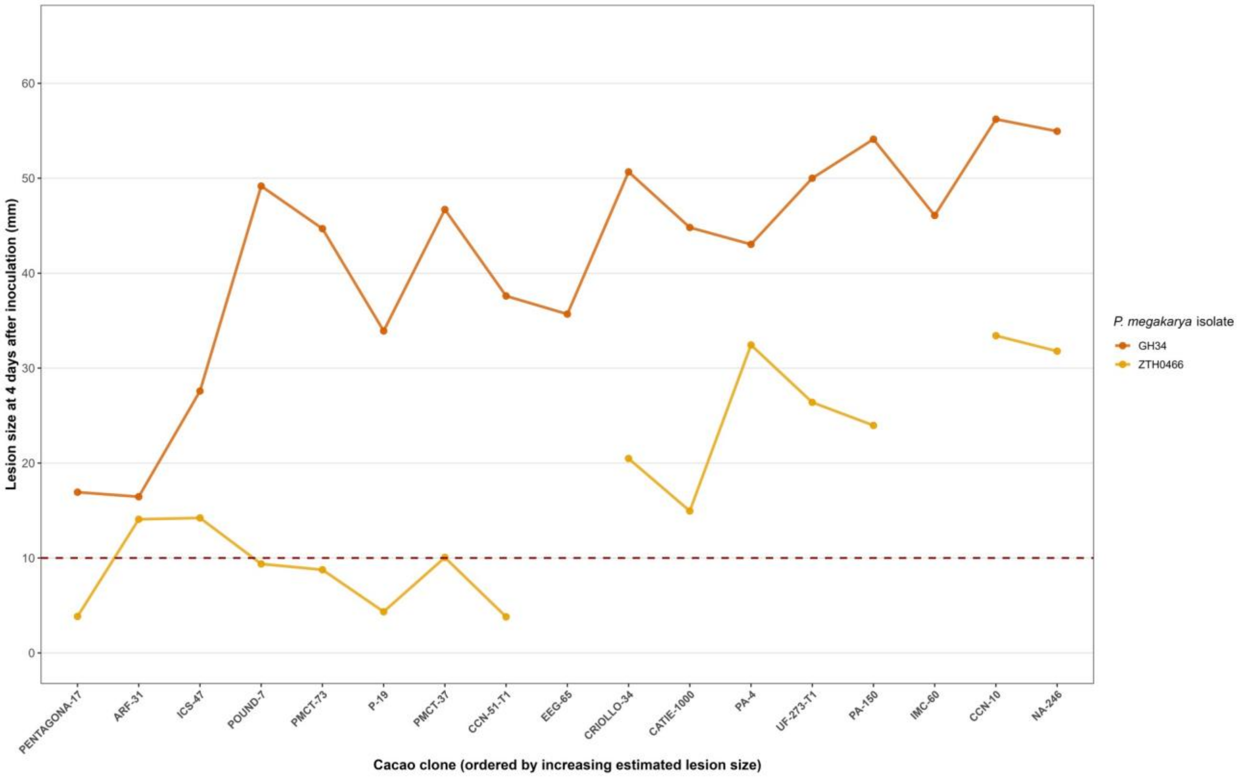


**Supplementary Fig. S3.** Intraspecific variation in estimated lesion size among *P. megakarya* isolates across cacao clones (conventions as in Fig. 3). These estimates are a subset of the fitted clone by isolate estimates from the primary lesion size model and are presented for visual comparison; no separate within species interaction was tested. The dashed line at 10 mm marks the approximate inoculation-patch width as a physical reference and was not used as a response-category threshold.

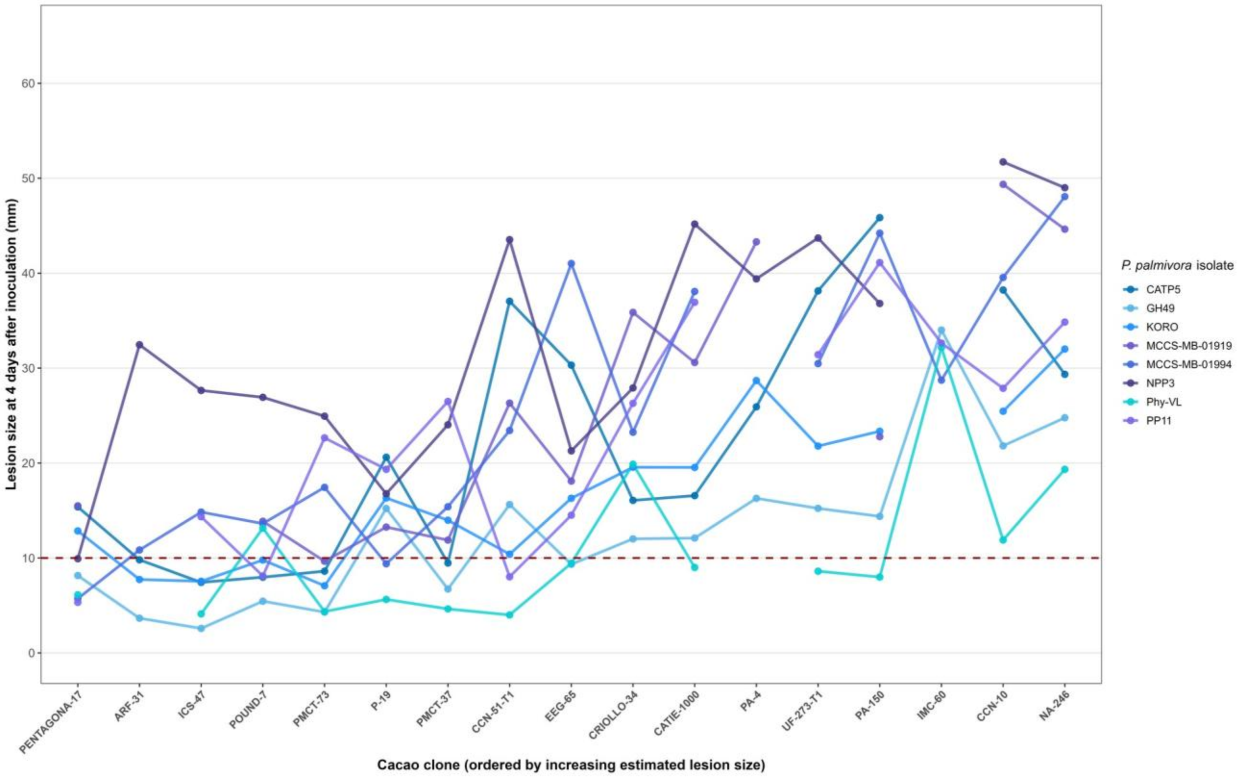


**Supplementary Fig. S4.** Intraspecific variation in estimated lesion size among *P. palmivora* isolates across cacao clones (conventions as in Fig. 3). These estimates are a subset of the fitted clone by isolate estimates from the primary lesion size model and are presented for visual comparison; no separate within species interaction was tested. The dashed line at 10 mm marks the approximate inoculation-patch width as a physical reference and was not used as a response-category threshold.

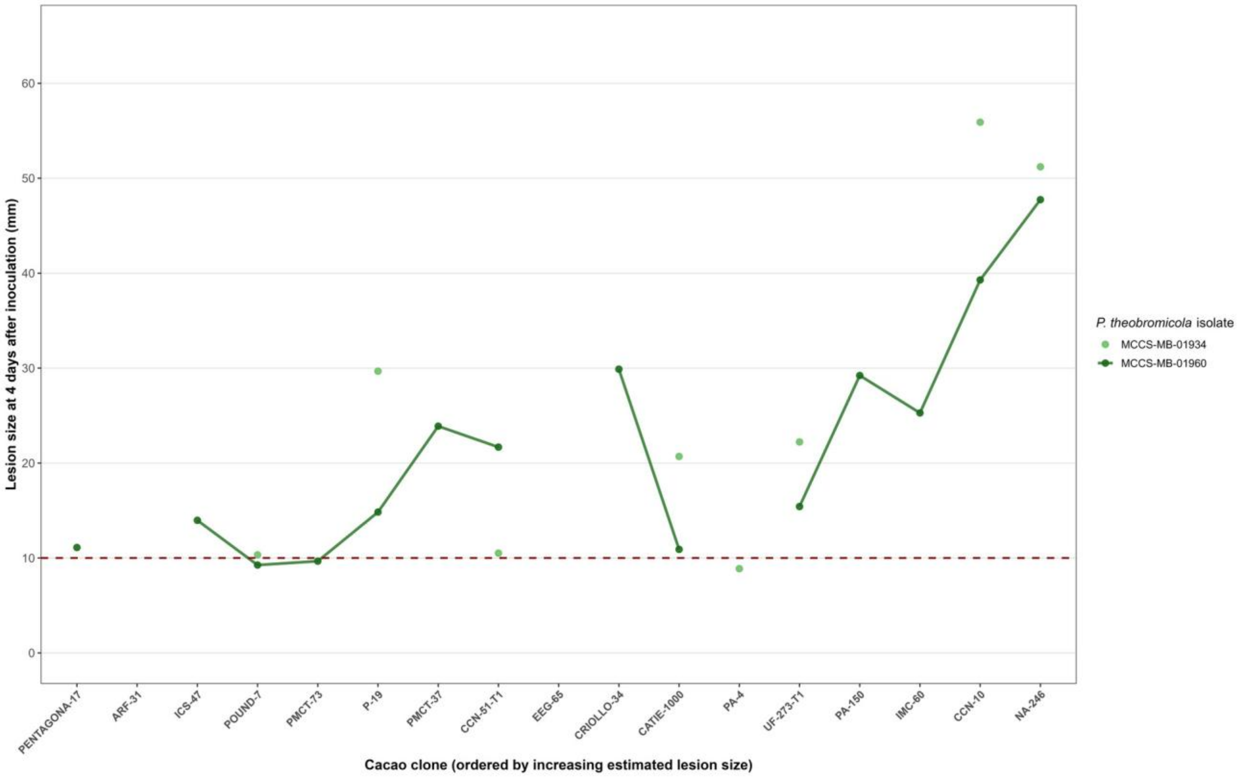


**Supplementary Fig. S5.** Intraspecific variation in estimated lesion size among *P. theobromicola* isolates across cacao clones (conventions as in Fig. 3). Points are shown for both retained isolates, but a line is drawn only for MCCS-MB-01960. MCCS-MB-01934 was represented for eight clones, and its estimates are displayed as unconnected points. These estimates are a subset of the fitted clone by isolate estimates from the primary lesion size model and are presented for visual comparison; no separate within species interaction was tested. The dashed line at 10 mm marks the approximate inoculation-patch width as a physical reference and was not used as a response-category threshold.

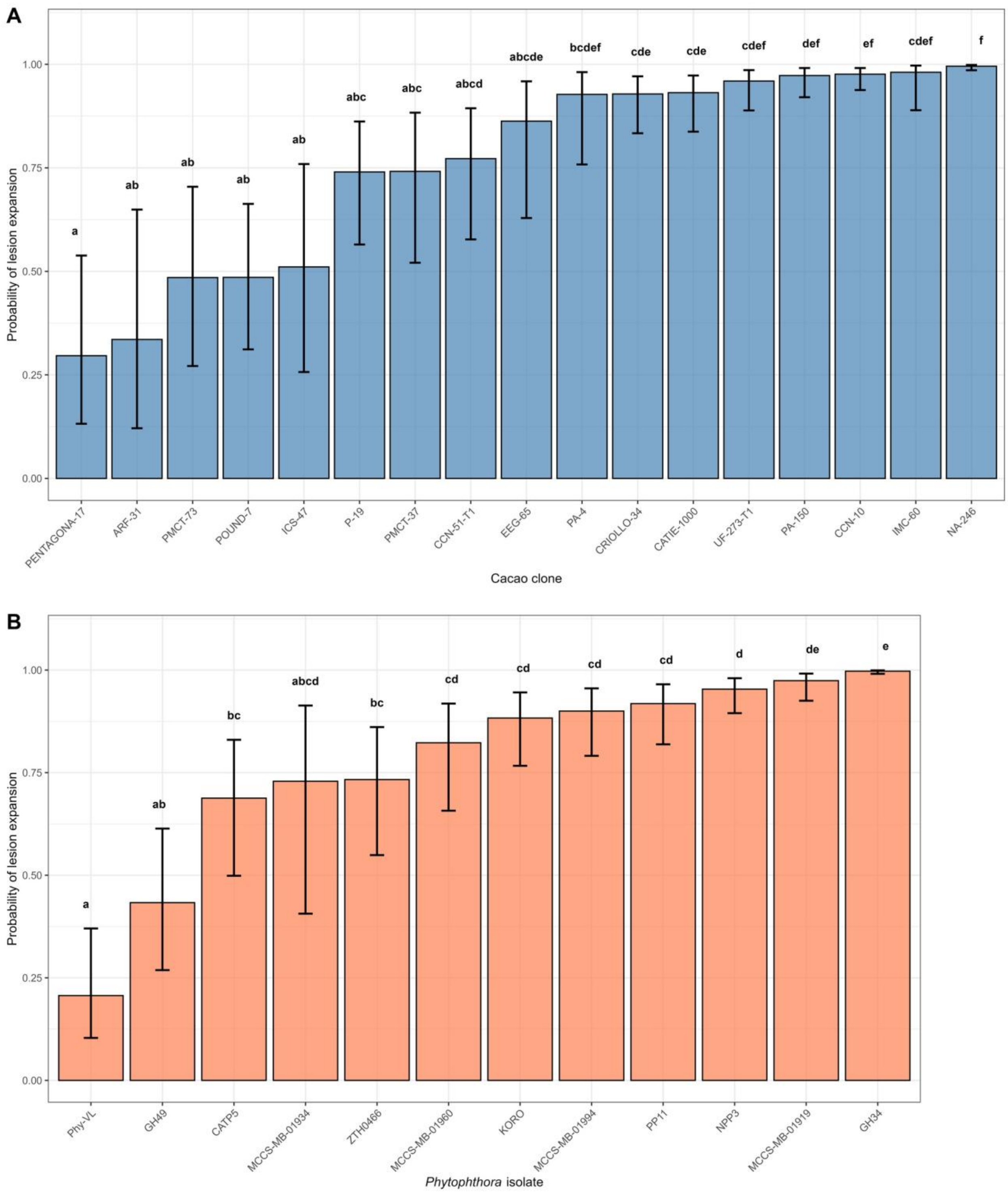


**Supplementary Fig. S6.** Model estimated probability of lesion expansion by cacao clone and *Phytophthora* isolate. Model estimated probabilities of lesion expansion by cacao clone (A) and *Phytophthora* isolate (B). Lesion expansion observed corresponds to reaction scores 2 and 3, whereas no lesion expansion detected corresponds to scores 0 and 1. Displayed values are estimated probabilities from the binomial generalized linear mixed model, and error bars are

95% confidence intervals back transformed from the logit scale. Lowercase letters denote comparisons adjusted using Tukey’s method on the logit scale at $\alpha = 0.05$. Shared letters indicate that a difference was not detected and do not demonstrate equivalence.
